# NMDAR Phosphoproteome Controls Synaptic Growth and Learning

**DOI:** 10.1101/2023.12.15.571360

**Authors:** Yasuhiro Funahashi, Rijwan Uddin Ahammad, Xinjian Zhang, Emran Hossen, Masahiro Kawatani, Shinichi Nakamuta, Akira Yoshimi, Minhua Wu, Huanhuan Wang, Mengya Wu, Xu Li, Md. Omar Faruk, Md Hasanuzzaman Shohag, You-Hsin Lin, Daisuke Tsuboi, Tomoki Nishioka, Keisuke Kuroda, Mutsuki Amano, Yukihiko Noda, Kiyofumi Yamada, Kenji Sakimura, Taku Nagai, Takayuki Yamashita, Shigeo Uchino, Kozo Kaibuchi

**Author notes:** These authors contributed contributed equally (Yasuhiro Funahashi, Rijwan Uddin Ahammad). Correspondence and proofs.

## Abstract

In the mammalian brain, NMDA receptors (NMDARs) activation triggers a calcium-dependent signal transduction cascade resulting in postsynaptic remodeling and behavioral learning. However, the phosphoprotein signal flow through this transduction network is poorly understood. Here, we show that NMDAR-dependent phosphorylation drives the assembly of protein signaling complexes that regulate synaptic morphology and behavior. We performed large-scale phosphoproteomic analyses of protein kinase target proteins in successive layers of the signaling network in mouse striatal/accumbal slices. NMDARs activation resulted in the phosphorylation of 194 proteins, including Rho GTPase regulators. CaMKII-mediated phosphorylation of ARHGEF2 increased its RhoGEF activity, thereby activating the RhoA-Rho-kinase pathway. Subsequent phosphoproteomics of Rho-kinase revealed 221 protein targets, including SHANK3. Experimental validation revealed a pathway from NMDAR-dependent calcium influx through CaMKII, ARHGEF2, Rho-kinase, and SHANK3 to coordinate assembly of an actin-tethered postsynaptic complex of SHANK3/NMDAR/PSD95/DLGAP3 for spine growth and aversive learning. These findings show that NMDARs initiate metabolic phosphorylation for learning.

## Introduction

Protein phosphorylation cascades regulate cellular dynamics. In the mammalian brain, N-methyl-D-aspartate receptor (NMDAR) activation triggers a calcium-dependent signal transduction cascade resulting in postsynaptic remodeling for learning and memory^1, 2, 3^. Important downstream effectors of NMDARs include Ca^2+^-dependent protein kinases, such as calcium/calmodulin-dependent protein kinase II (CaMKII), that phosphorylate and regulate various target proteins such as signal-related proteins, ion channel proteins, and transcription factors^4, 5^. However, progress in this field has been slowed in part due to the lack of tools to identify protein kinase phosphorylation targets at a large scale. Although several CaMKII target proteins have been identified, such as NR2B (a subunit of NMDARs) and GluR1 (a subunit of α-amino-3-hydroxy-5-methyl-4-isoxazole propionic acid receptors (AMPARs))^6, 7, 8^, research on the overall signal flow controlled by phosphorylation in this historically important cell pathway remains immature.

The nucleus accumbens (NAc) of the basal ganglia plays a pivotal role in reward and aversive learning^9, 10^. Approximately, 95% of all neurons in the NAc are medium spiny neurons (MSNs). MSNs are subdivided into dopamine receptor D1-expressing MSNs (D1R-MSNs) and dopamine receptor D2-expressing MSNs (D2R-MSNs), and they are primarily involved in reward and aversive learning, respectively^10^. The activities of D1R-MSNs and D2R-MSNs are regulated by various neuromodulators including dopamine, acetylcholine, and adenosine and neurotransmitters including glutamate^11, 12^. We previously developed a kinase-oriented phosphoproteomic method named Kinase-Oriented Substrate Screening (KIOSS), which uses affinity beads coated with pS/T-binding domains, including the tryptophan–tryptophan (WW) and forkhead associated (FHA) domains and 14-3-3 proteins to enrich phosphorylated proteins^13, 14, 15^. Using KIOSS, we identified over 100 candidate substrates of PKA (including the Rap1 GTPase regulators, RASGRP2 and Rap1GAP) and mitogen-activated protein kinase (MAPK) (including KCNQ2, a subunit of voltage-gated K^+^ channels) downstream of dopamine in the mouse NAc^13^. In a subsequent study, we found that PKA stimulated MAPK via Rap1, thereby promoting neuronal excitability for reward learning through the phosphorylation of KCNQ2^16^.

Here, we performed the first phosphoproteomic analysis of the NMDAR pathway in the striatum/NAc. We identified 194 proteins whose phosphorylation was promoted by NMDA and demonstrated that NMDA induced activation of the CaMKII-ARHGEF2-Rho-kinase pathway followed by phosphorylation of 221 Rho-kinase target proteins including SHANK3. Finally, we demonstrate that our identified NMDAR signal transduction pathway results in the assembly of protein signaling complexes linking membrane receptors with kinase regulators and adaptors for aversive learning in D2R-MSN of the NAc. Our high-throughput phosphoproteomic methods provide a tool to unlock major conceptual advances in various biological signaling cascades hitherto impossible in conventional studies.

## Results

### Phosphoproteomic analysis of NMDAR signaling

To investigate novel phosphorylation events downstream of NMDAR signaling, we utilized acute slices of mouse striatum/NAc and treated them with either high potassium (high K^+^) or an NMDAR agonist (NMDA) to induce Ca^2+^ influx. Neuronal Ca^2+^ influx activates CaMKII and enhances the phosphorylation of known CaMKII substrates, including NR2B and GluR1^6, 7, 8^. We first confirmed the phosphorylation of these proteins by immunoblot analysis after the stimulation of striatal/accumbal slices with high K^+^ or NMDA. Both high K^+^ and NMDA stimulation increased the phosphorylation not only of NR2B at S1303 and of GluR1 at S831, but also of CaMKII at T286, an autophosphorylation site of CaMKII and an indicator of CaMKII activation (Fig. 1a, b). These phosphorylation levels peaked between 15 and 30 seconds (s) after high K^+^ and NMDA stimulation and returned to basal levels within 5 minutes (min) (Fig. 1a, b).

**Fig. 1:**
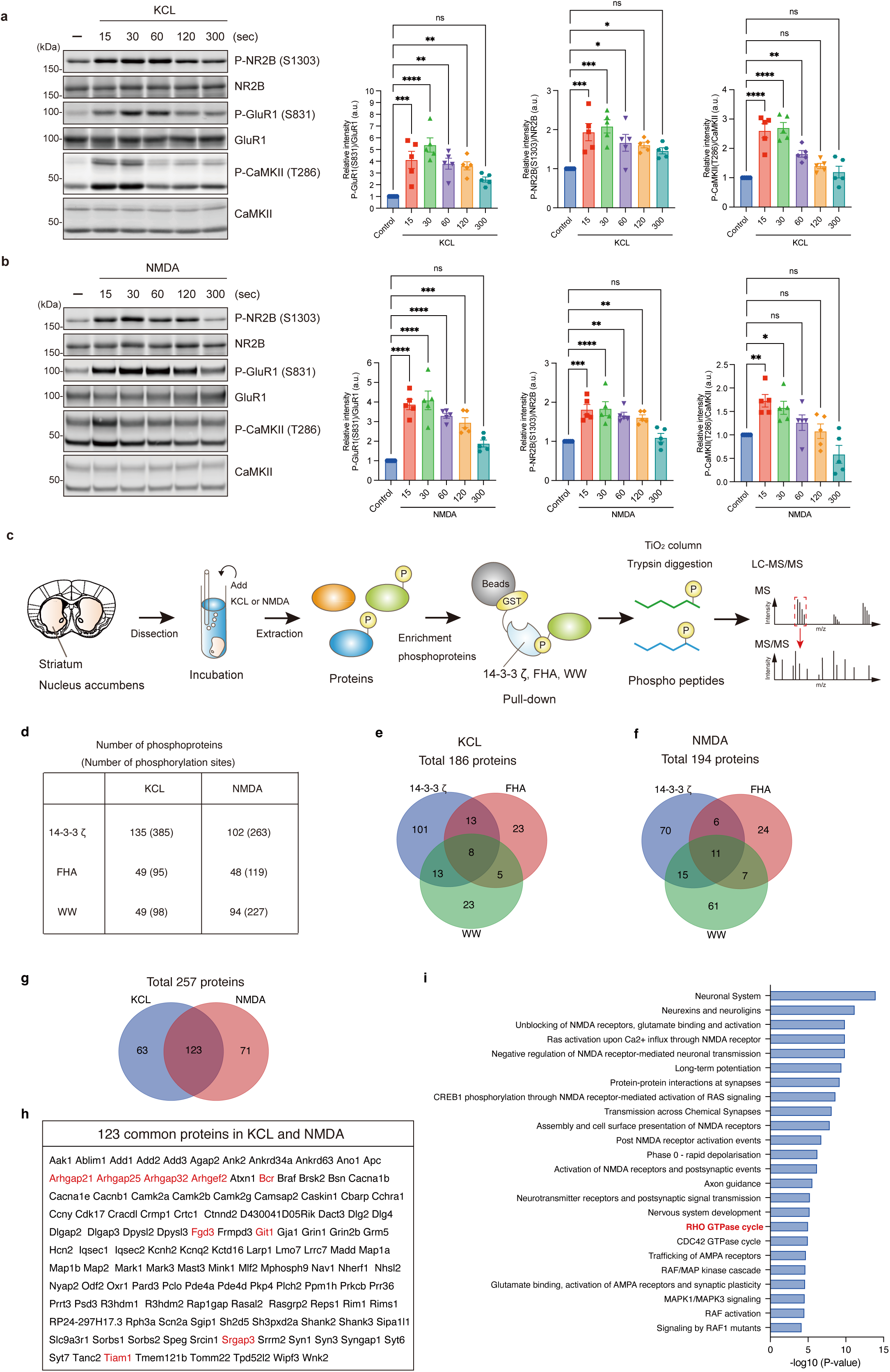
Phosphoproteomic analysis of NMDAR signaling. **(a)** High K^+^ induced the phosphorylation of NR2B (S1303), GluR1 (S831), and CaMKII (T286). Striatal/accumbal slices were treated with KCL (40 mM) for 15, 30, 60, 120, or 300 s and subjected to immunoblot analysis. Data present the mean ± SEM of five independent experiments. One-way ANOVA followed by Dunnett’s multiple comparisons test. *p<0.05, **p<0.01, ***p<0.001, ****p<0.0001. ns: not significant. **(b)** NMDA induced the phosphorylation of NR2B (S1303), GluR1 (S831), and CaMKII (T286). Striatal/accumbal slices were treated with NMDA (100 µM) for 15, 30, 60, 120, or 300 s and subjected to immunoblot analysis. Data present the mean ± SEM of five independent experiments. One-way ANOVA followed by Dunnett’s multiple comparisons test. *p<0.05, **p<0.01, ***p<0.001, ****p<0.0001. ns: not significant. **(c)** Schematic representation of the Kinase-Oriented Substrate Screening (KIOSS) method to identify phosphoproteins associated with Ca^2+^ influx using high K^+^ or NMDA in striatal/accumbal slices. Extracts of these striatal/accumbal slices were applied to affinity beads coated with pS/T-binding domains (14-3-3 ζ, FHA, and WW) to enrich the phosphoproteins. The bound proteins were digested using trypsin and subjected to LC-MS/MS to identify the phosphorylated proteins and their phosphorylation sites. **(d)** The number of phosphoproteins bound to each of the WW, FHA, and 14-3-3ζ baits after KCL and NMDA treatment is shown. **(e-f)** The number of phosphoproteins bound to each bait is compared using a Venn diagram. **(g)** Venn diagrams showing the number of identified phosphoproteins identified by KCL and NMDA. **(h)** List of common proteins identified by KCL and NMDA. **(i)** Pathway analysis using the Reactome database (available online at: http://www.reactome.org) of 123 proteins commonly identified by KCL and NMDA.

Phosphoproteomic analysis was then performed using the KIOSS method to identify phosphoproteins associated with Ca^2+^ influx (Fig. 1c). Striatal/accumbal slices were treated with high K^+^ or NMDA for 15 s, and extracts were then applied to affinity beads coated with pS/T-binding modules, including 14-3-3ζ protein, CHECK2-FHA domain, and PIN1-WW domain, to enrich the phosphorylated proteins. The binding motif preferences of the aforementioned proteins for phosphoproteins varied, with 14-3-3ζ preferring the [(R/K)XX (pS/pT)] motif, FHA preferring the [(pS/pT)P] motif with acidic amino acid at +3 position and basic amino acids at −3 position, and WW preferring the proline-rich (pS/pT)P motif^13, 15, 17^. Proteins bound to 14-3-3ζ, FHA domain, or WW domain were digested with trypsin/Lys-C and loaded onto a TiO_2_ column to enrich phosphopeptides. This was followed by liquid chromatography/tandem mass spectrometry (LC-MS/MS) to identify phosphorylated proteins and their phosphorylation sites. The high K^+^ treatment identified 135, 49, and 49 phosphoproteins from 14-3-3ζ, FHA, and WW, respectively (Fig. 1d), while the NMDA treatment identified 102, 48, and 94 phosphoproteins from each bait, respectively (Fig. 1d). Venn diagrams showed that the high K^+^ and NMDA treatments identified 186 and 194 non-overlapping phosphoproteins in each bait, respectively (Fig. 1e, f). A total of 257 proteins were identified, of which 123 (48%) were commonly stimulated by high K^+^ and NMDA (Fig. 1g, h). The phosphorylated proteins and their phosphorylation sites that were increased more than 2-fold compared to those in the control and were detected at least twice in three or more independent experiments are summarized in Table 1. Detailed information about the phosphorylation sites is summarized in Supplementary Table 1 and can also be found in a Kinase-Associated Neural Phospho-Signaling (KANPHOS) database (https://kanphos.neuroinf.jp), which we developed to provide information about the phosphorylation signals identified by our methods as well as those reported previously^18^. An analysis of the identified phosphopeptide sequence revealed that approximately 42% of the phosphorylation sites contained basic amino acids (arginine or lysine) at the −3 position [(R/K)XX(pS/pT)] or the −2 position [(R/K)X(pS/pT)], consistent with the consensus motif for CaMKII or PKC^19^. This suggests that CaMKII, PKC, or a related kinase is responsible for these phosphorylation events (Supplementary Fig. 1a, b). Additionally, approximately 22% of the phosphorylation sites contained proline at the +1 position [(pS/pT)P], consistent with the consensus motif for MAPK1/3^19^. This suggests that MAPK1/3 or other proline-oriented kinases are responsible for these phosphorylation events (Supplementary Fig. 1a, b). This finding provides further evidence that MAPK1/3 is activated downstream of CaMKII or PKC^20^. Furthermore, Approximately 45% of the phosphorylation sites did not match any of the CaMKII, PKC, or MAPK1/3 motifs, suggesting that these are phosphorylation sites for other kinases (Supplementary Fig. 1a, b). Notably, most of these proteins and their phosphorylation sites have not yet been reported as a part of the NMDAR signaling pathway.

**Table 1.**
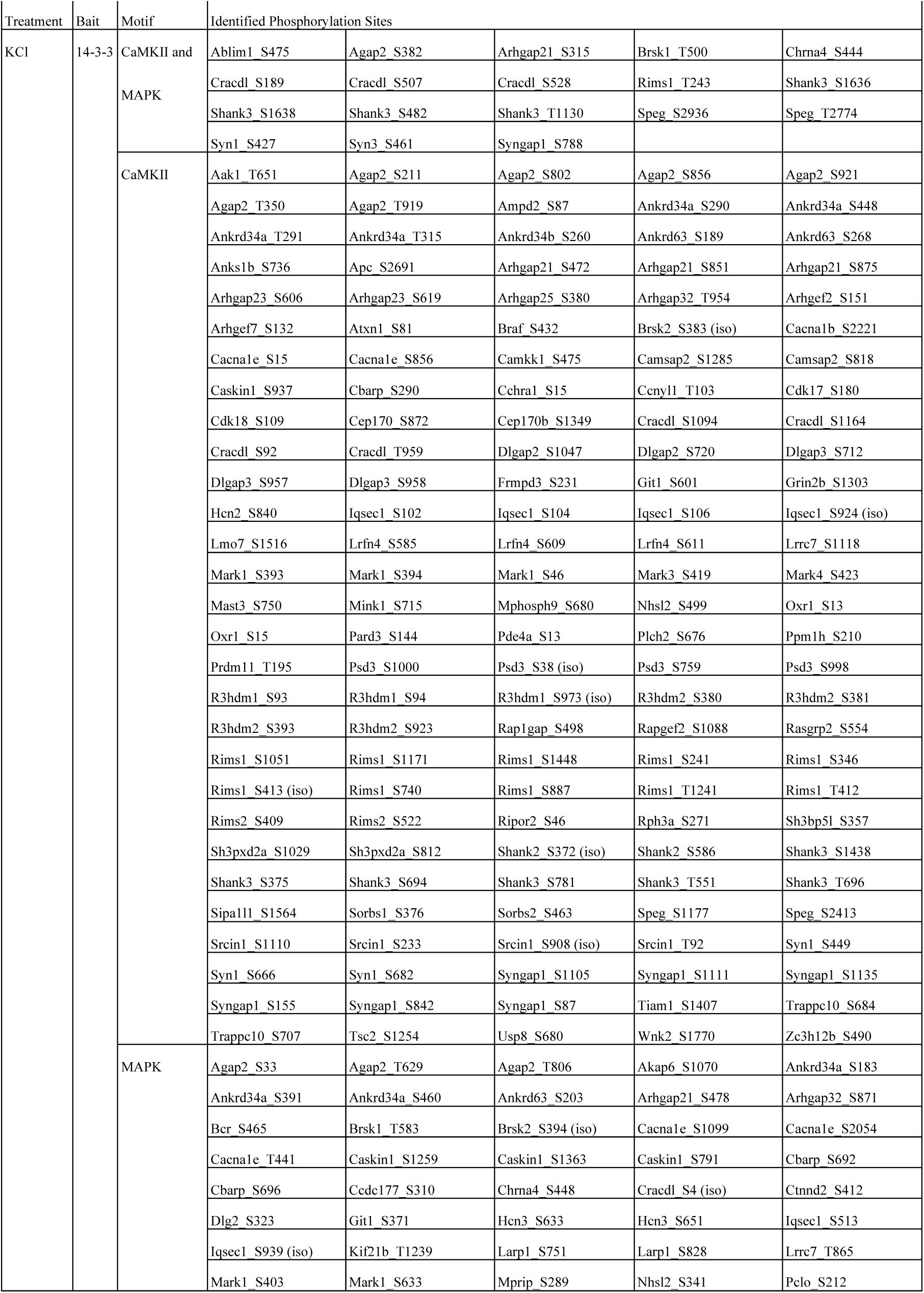
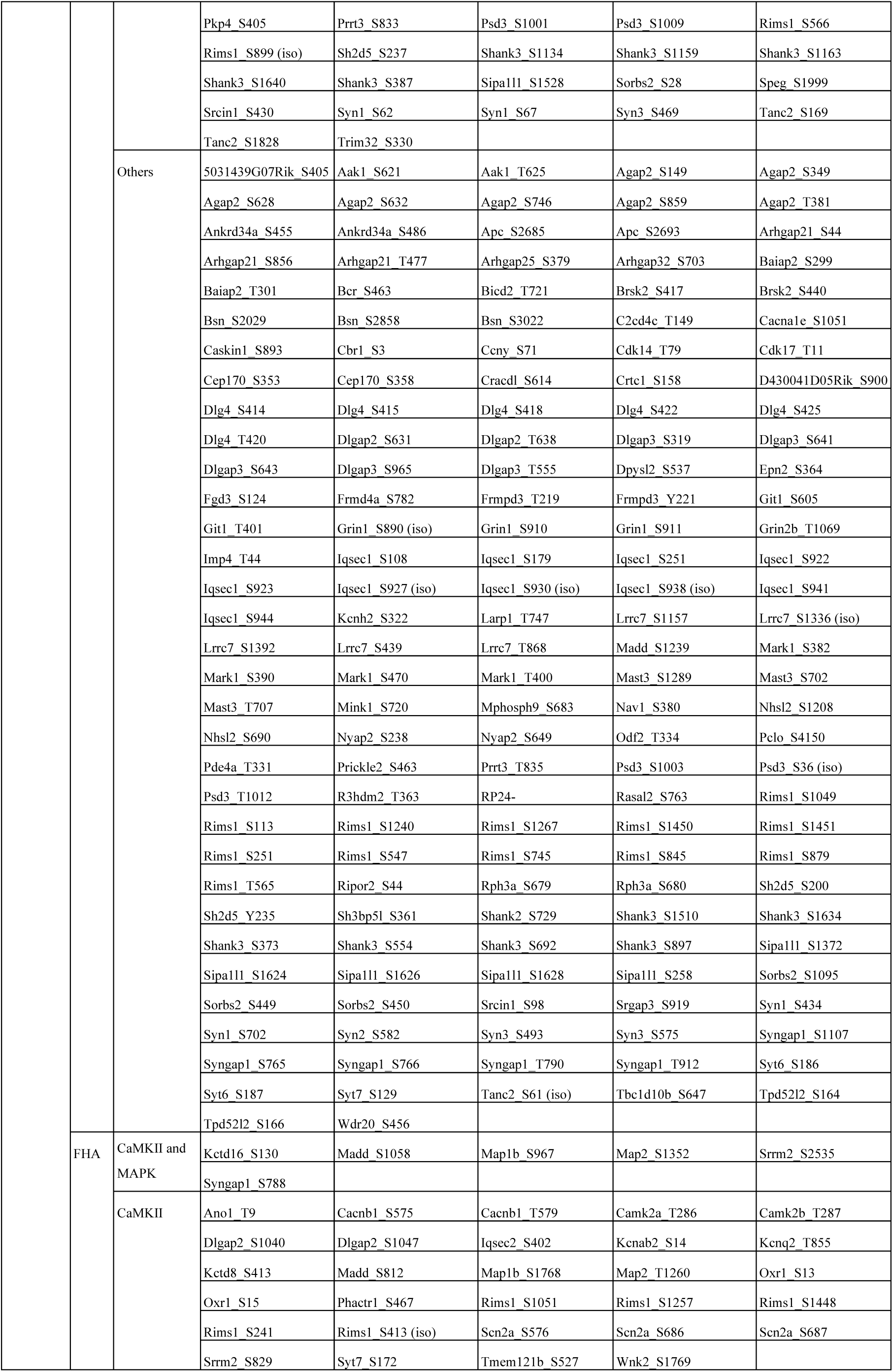
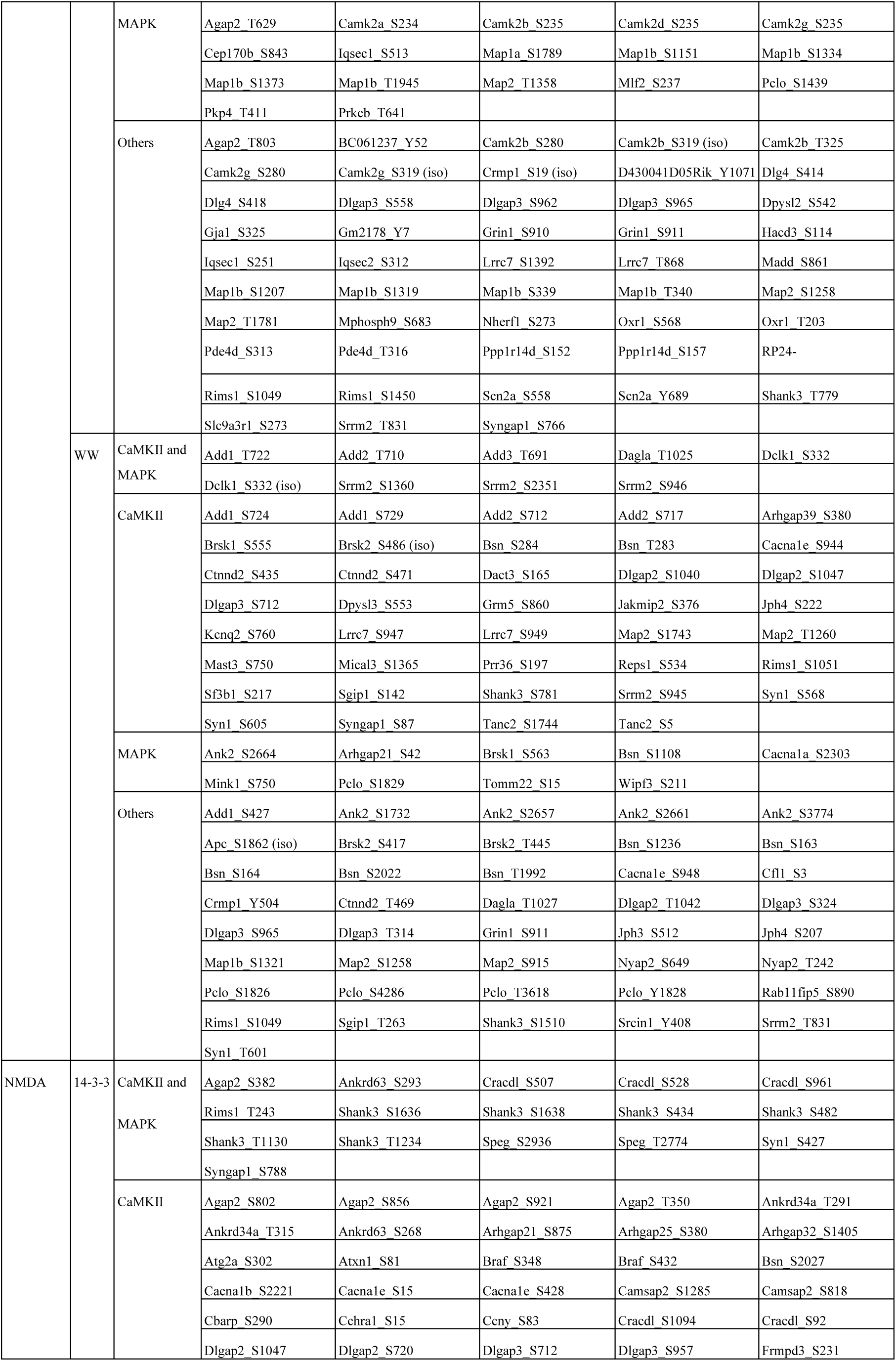
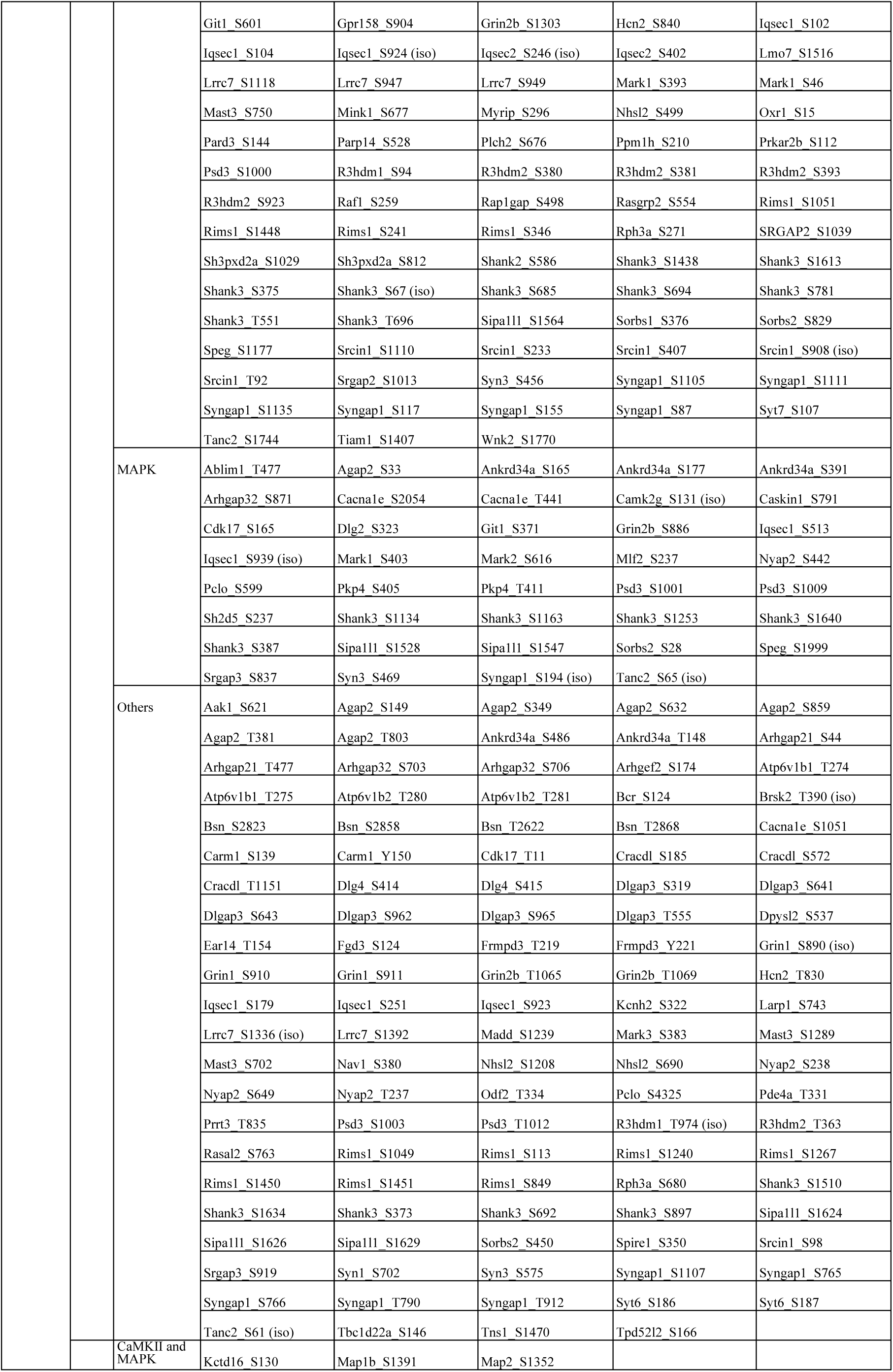
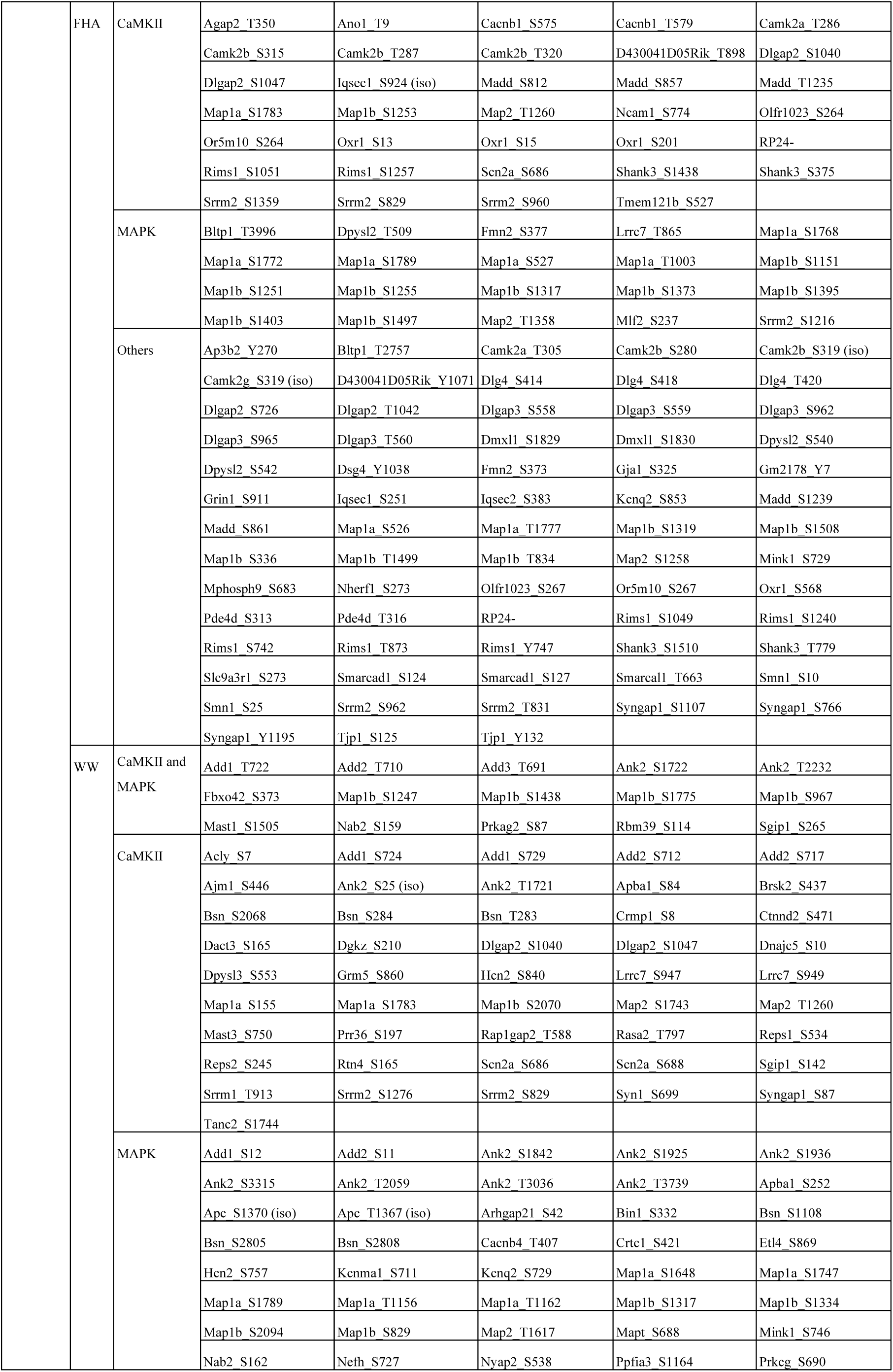
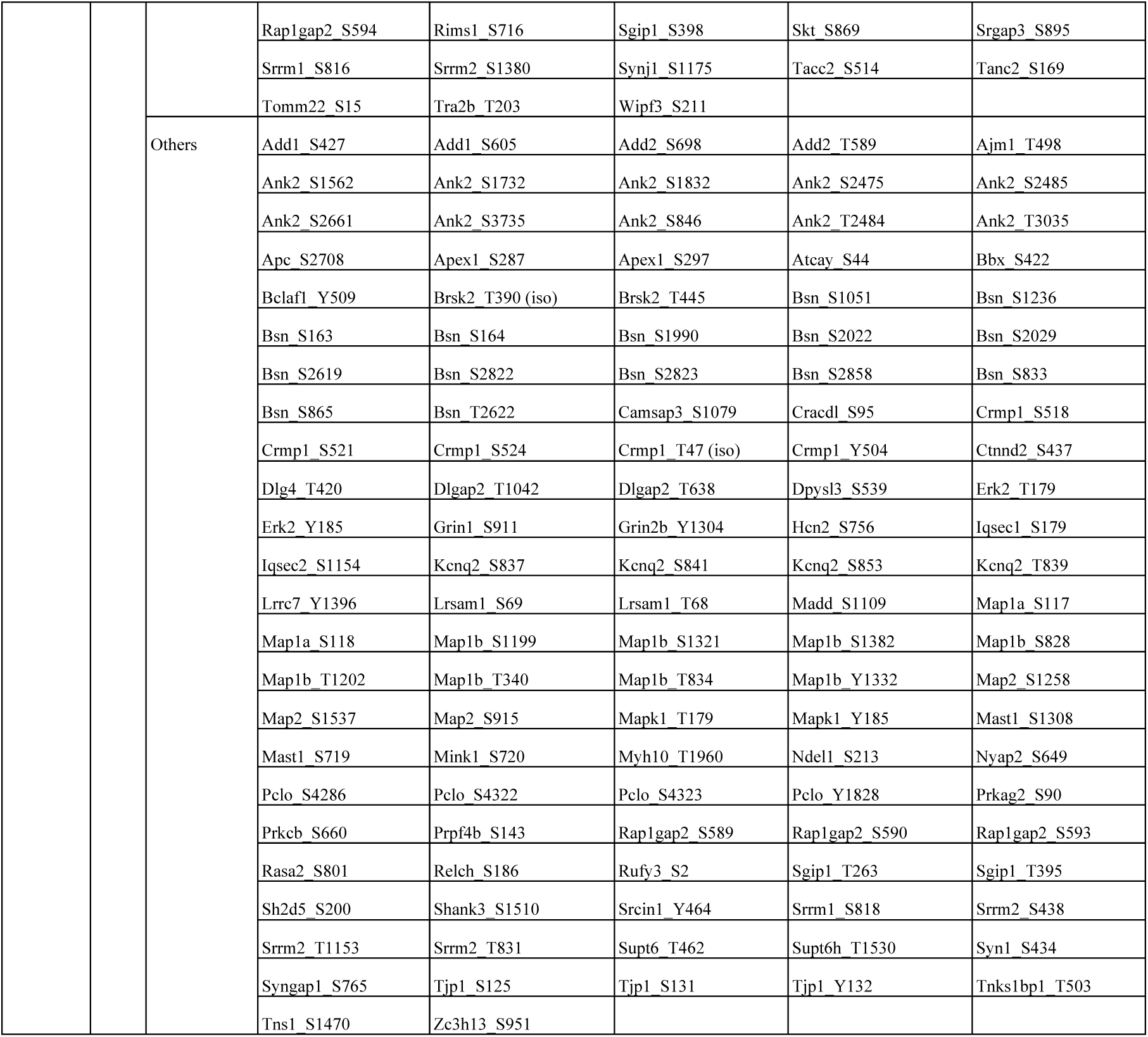
Protein phosphorylation induced by high K^+^ and NMDA in the mouse striatum. Striatal slices were treated with KCL or NMDA to induce CaMKII-mediated phosphorylation. The extracts of these striatal slices were then applied to affinity beads coated with 14-3-3 ζ, FHA, or WW to enrich the phosphorylated proteins. The bound proteins were digested using trypsin and subjected to LC-MS/MS to identify the phosphorylated proteins and their phosphorylation sites. The reproducible phosphorylation sites were stimulated by more than 2-fold higher than those of the control and at least twice in more than three independent experiments (14-3-3 ζ, n=5; FHA, n=3; WW, n=3). The results are summarized. See also Supplementary Table 1.

To identify the signaling pathways associated with the proteins obtained upon stimulation with high K^+^ and NMDA, we utilized a curated pathway database, Reactome (http://www.reactome.org/), and entered 123 proteins that were commonly stimulated by high K^+^ and NMDA. Pathway analysis revealed that ‘Rho GTPase cycle’ was one of the NMDA-related pathways (Fig. 1i). Several Rho family GTPase regulators, including the Rho family GEFs (*Arhgef2, Fgd3,* and *Tiam1*) and Rho family GAPs (*Arhgap21*, *Arhgap25*, *Arhgap32*, *Bcr*, *Git1*, *and Srgap3*), were included among the proteins associated with ‘the Rho GTPase cycle’ (Fig. 1h). As RhoA is involved in synaptic plasticity and its activation depends on CaMKII^21^, we focused on RhoA-specific regulators, including ARHGEF2 (also known as GEF-H1) and ARHGAP21 (also known as ARHGAP10, KIAA1424)^22, 23^.

### CaMKII phosphorylates ARHGEF2 and ARHGAP21, and activates RhoA–Rho-kinase pathway downstream of NMDARs

We initially focused on ARHGEF2, which plays a crucial role in regulating the spatiotemporal activation of RhoA by converting RhoA from its inactive GDP-bound form to its active GTP-bound form^24^. CaMKII phosphorylated GST-ARHGEF2 (84-130 aa) *in vitro*, with the level of phosphorylation being approximately 75% lower in the T103A mutant (Fig. 2a). To further confirm this finding, we generated a phosphospecific antibody for the ARHGEF2 phosphorylated at T103 and verified that this antibody specifically recognized phosphorylated ARHGEF2 by CaMKII, but did not cross-react with non-phosphorylated ARHGEF2 (Supplementary Fig. 2a). Phosphorylation of ARHGEF2 at T103 was increased by co-expression of the constitutively active form of CaMKIIα (CaMKIIα-CA) in COS7 cells, but not by that of CaMKIIα-WT or the kinase-dead form of CaMKIIα (CaMKIIα-CATKD) (Fig. 2b). These results indicate that CaMKII directly phosphorylates ARHGEF2 at T103.

**Fig. 2:**
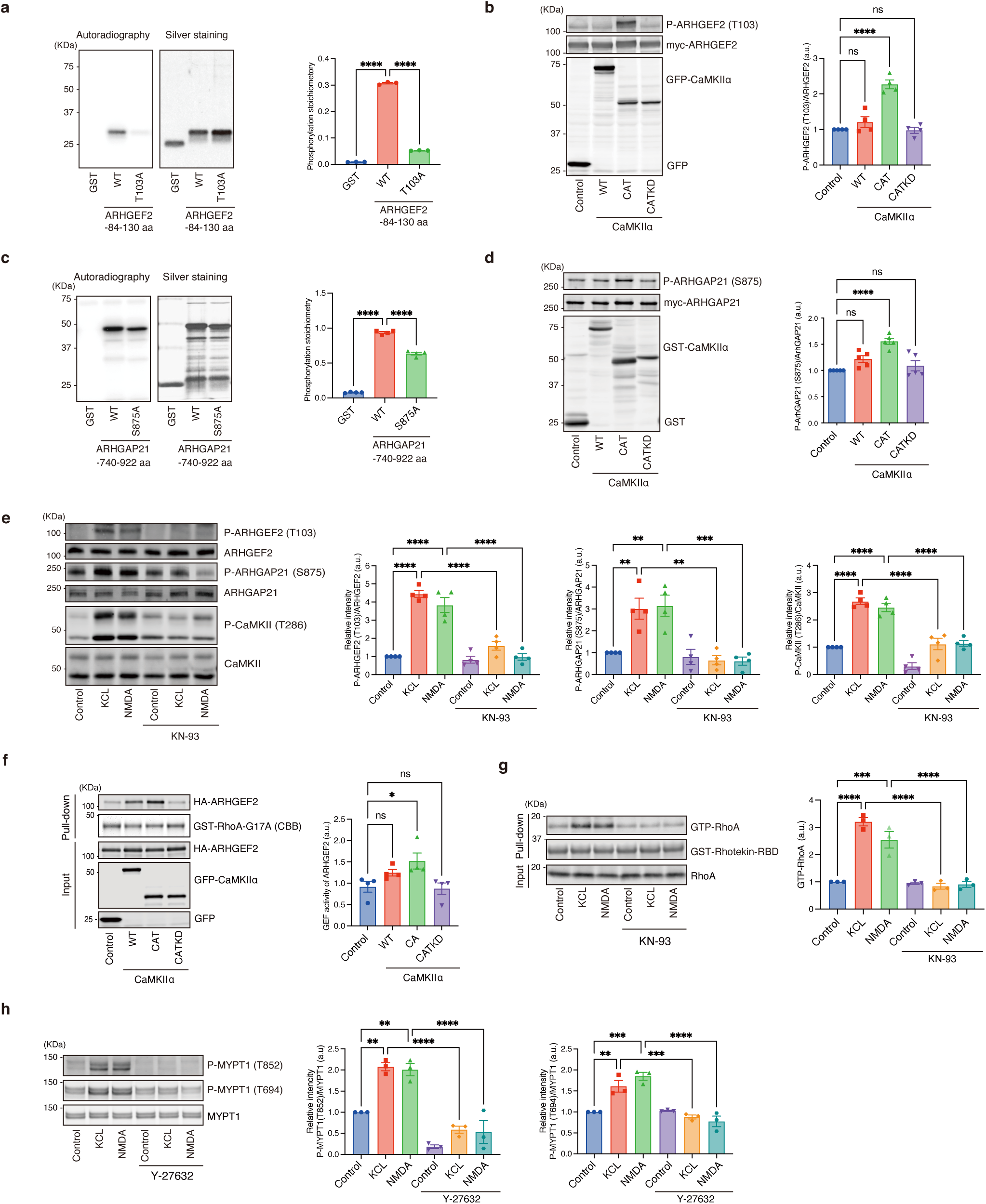
CaMKII phosphorylates ARHGEF2 and ARHGAP21, and activates RhoA–Rho-kinase pathway downstream of NMDARs **(a)** CaMKII phosphorylated ARHGEF2 (T103) *in vitro*. Data represent the mean ± SEM of three independent experiments. One-way ANOVA followed by Tukey’s multiple comparisons test. ****p<0.0001. **(b)** CaMKII phosphorylated ARHGEF2 (T103) in COS7 Cells. Data represent the mean ± SEM of four independent experiments. One-way ANOVA followed by Dunnett’s multiple comparisons test. ****p<0.0001. **(c)** CaMKII phosphorylated ARHGAP21 (S875) *in vitro*. Data represent the mean ± SEM of four independent experiments. One-way ANOVA followed by Tukey’s multiple comparisons test. ****p<0.0001. **(d)** CaMKII phosphorylated ARHGAP21 (S875) in COS7 Cells. Data represent the mean ± SEM of five independent experiments. One-way ANOVA followed by Dunnett’s multiple comparisons test. ****p<0.0001. **(e)** CaMKII phosphorylated ARHGEF2 (T103) and ARHGAP21 (S875) downstream of NMDARs in the striatum/NAc. Striatal/accumbal slices were pretreated with KN-93 (50 µM) for 4 h and subsequently treated with KCL (40 mM) or NMDA (100 µM) for 15 s and subjected to immunoblot analysis. Data present the mean ± SEM of four independent experiments. Two-way ANOVA followed by Tukey’s multiple comparisons test. **p<0.01, ***p<0.001, ****p<0.0001. **(f)** CaMKII-CAT increased the phosphorylation of ARHGEF2 (T103) and the guanine nucleotide exchange activity of ARHGEF2 in COS7 cells. The guanine nucleotide exchange activity of ARHGEF2 was measured by GST-RhoA-G17A pulldown assay. Data represent the mean ± SEM of four independent experiments. One-way ANOVA followed by Tukey’s multiple comparisons test. *p<0.05, ****p<0.0001. ns: not significant. **(g)** High K^+^ and NMDA induced RhoA activity in the striatum/NAc. Striatal/accumbal slices were pre-treated with KN-93 (50 µM) for 4 h and subsequently treated with KCL (40 mM) or NMDA (100 µM) for 15 s. The lysates were incubated with GST-Rhotekin-RBD. Data represent the mean ± SEM of three independent experiments. Two-way ANOVA followed by Tukey’s multiple comparisons test. ***p<0.001, ****p<0.0001. **(h)** High K^+^ and NMDA induced phosphorylation of MYPT1 at T852 and T694. Striatal/accumbal slices were pretreated with Y-27632 (20 µM) for 1 h, treated with KCL (40 mM) or NMDA (100 µM) for 15 s, and subjected to immunoblot analysis. Data represent the mean ± SEM of three independent experiments. Two-way ANOVA followed by Tukey’s multiple comparisons test. **p<0.01, ***p<0.001, ****p<0.0001.

On the other hand, RhoGAPs are critical for the hydrolysis of GTP bound to Rho, Rac, and/or Cdc42, thereby inactivating the Rho family GTPases^25^. Among the identified RhoGAPs, ARHGAP21 is associated with RhoA^22^. CaMKII phosphorylated GST-ARHGAP21 (740-922 aa) *in vitro*, with the level of phosphorylation being approximately 40% lower in the S875A mutant (Fig. 2c). We generated a phosphospecific antibody for the ARHGAP21 phosphorylated at T875 and verified that this antibody specifically recognized CaMKII-phosphorylated ARHGAP21 but did not cross-react with non-phosphorylated ARHGAP21 (Supplementary Fig. 2b). Phosphorylation of ARHGAP21 at S875 was increased by the co-expression of CaMKII-CA in COS7 cells, but not by that of CaMKII-WT or CaMKII-CATKD (Fig. 2d). These results indicate that CaMKII directly phosphorylates ARHGAP21 at S875.

Furthermore, we investigated whether CaMKII phosphorylated ARHGEF2 and ARHGAP21 in the striatum/NAc. Both high K^+^ and NMDA increased the phosphorylation of ARHGEF2 at T103 and ARHGAP21 at S875, which was inhibited by pretreatment with a CaMKII inhibitor (KN-93) (Fig. 2e). These results suggest that CaMKII phosphorylates ARHGEF2 and ARHGAP21 downstream of NMDARs in the striatum/NAc.

Considering that RhoGEFs play a more critical role in RhoA activation than RhoGAPs^26, 27^, we analyzed ARHGEF2 phosphorylation in detail. To evaluate whether the phosphorylation of ARHGEF2 affects its guanine nucleotide exchange activity, the guanine nucleotide exchange activity of ARHGEF2 was measured using an affinity precipitation assay with GST-RhoA-G17A (a nucleotide-free mutant of RhoA that preferentially interacts with activated RhoA GEF)^28^. Co-expression of ARHGEF2 with CaMKII-CA increased the amount of ARHGEF2 precipitated with GST-RhoA-G17A (Fig. 2f), suggesting that CaMKII phosphorylates ARHGEF2, thereby activating GEF activity on RhoA. To investigate whether CaMKII affects RhoA-GTP levels in the striatal/accumbal slices, we used beads coated with Rhotekin-RBD (a RhoA effector) and performed an affinity precipitation assay to measure RhoA-GTP levels. Treatment of striatal/accumbal slices with high K^+^ or NMDA increased RhoA-GTP levels, whereas pretreatment with a CaMKII inhibitor (KN-93) prevented the high K^+^- or NMDA-induced increase in RhoA-GTP levels (Fig. 2g).

Previous pharmacological studies have suggested that Rho-kinase/ROCK (another RhoA effector) is required for the NMDA-induced structural plasticity of dendritic spines^21^. Therefore, we investigated whether high K^+^ or NMDA regulates the activity of Rho-kinase. Treatment of striatal/accumbal slices with high K^+^ or NMDA induced the phosphorylation of myosin phosphatase-targeting subunit 1 (MYPT1) at T852 and T694 (Fig. 2h), which is the best known substrate of Rho-kinase^29^. Pretreatment with a Rho-kinase inhibitor (Y-27632) inhibited the high K^+^- or NMDA-induced phosphorylation of MYPT1 at T852 and T694 (Fig. 2h). These results indicate that NMDA activates the RhoA-Rho-kinase pathway via the CaMKII-mediated phosphorylation of Rho GTPase regulators, including ARHGEF2.

### Aversive stimuli induce CaMKII–RhoA–Rho-kinase pathway in D2R-MSNs of the NAc

Prominent glutamate input to the NAc originates from the ventral hippocampus, basolateral amygdala, and prefrontal cortex^30^. Acute foot shock stress increases glutamate release from the prefrontal cortex, and excess glutamate allows neuronal Ca^2+^ influx, thereby activating CaMKII^31^. Local glutamatergic transmission in the NAc plays a role in aversive learning^32^, and D2R-MSNs in the NAc are known to be involved in aversive behaviors^10^. Therefore, we first examined the real-time activity of D2R-MSNs in the NAc during aversive stimuli. AAV-Flex-GCaMP6f was injected into the NAc of adenosine A2a receptor (Adora2a)-Cre transgenic mice^33, 34^ to specifically express GCaMP6f in D2R-MSNs. Thereafter, the mice were subjected to electric foot shock, and Ca^2+^ signaling was recorded transiently through wireless fiber photometry (Fig. 3a). Electric foot shock immediately and transiently increased Ca^2+^ influx in D2R-MSNs, which was suppressed by the subcutaneous injection of an NMDAR antagonist (MK-801) (Fig. 3b-d). Based on these results, we investigated the relationship between aversive stimuli and the CaMKII-RhoA-Rho-kinase pathway in the NAc.

**Fig. 3:**
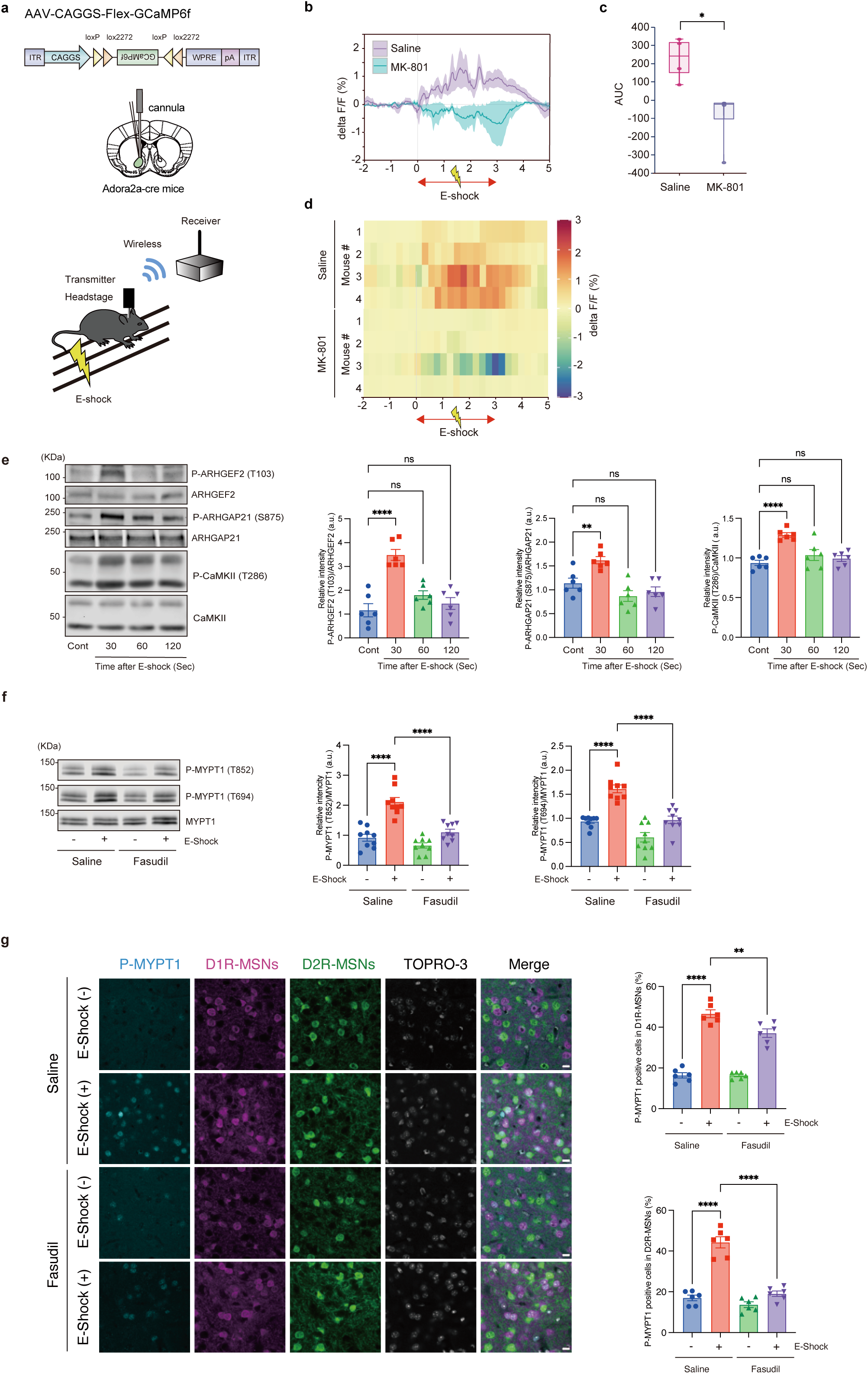
Aversive stimuli induce CaMKII–RhoA–Rho-kinase pathway in D2R-MSNs of the NAc. **(a)** AAV-CAGGS-Flex-GCaMP6f was injected into the NAc of Adora2a-Cre mice. Calcium activity during electric foot shock was continuously recorded using wireless fiber photometry. **(b)** The average Ca^2+^ trace from 2 s before to 5 s after foot shock. **(c)** Quantification of area under curve (AUC) of ΔF/F GCaMP6f signal. Data present the mean ± SEM (n=4 mice per each condition). Mann-Whitney U test. *p<0.05. **(d)** The heatmap shows the Ca^2+^ responses of GCaMP6-expressing D2R-MSNs before and during electric foot shock among individual animals in the saline and MK-801 groups. **(e)** Electric foot shock (0.7 mA, 3 s) increased the phosphorylation of ARHGEF2 (T103), ARHGAP21 (S875), and CaMKII (T286) in the NAc. Data present the mean ± SEM (n=6 mice per each condition). One-way ANOVA followed by Dunnett’s multiple comparisons test. **p<0.01, ****p<0.0001. ns: not significant. **(f)** Rho-kinase inhibitor (20 mg/kg fasudil, intraperitoneally) inhibited the electric foot shock-induced the phosphorylation of MYPT1 at T694 and T852. Data present the mean ± SEM (n=9 mice per each condition). Two-way ANOVA followed by Tukey’s multiple comparisons test. ****p<0.0001. **(g)** Electric foot shock (0.7 mA, 3 s) increases the number of phospho-MYPT1 (T694)-positive MSNs in the NAc of D1R-tdTomato/D2R-mVenus mice. Data present the mean ± SEM (n=6 mice per each condition). Two-way ANOVA followed by Tukey’s multiple comparisons test. **p<0.01, ****p<0.0001.

To examine whether electric foot shock activated the CaMKII-RhoA-Rho-kinase pathway in the NAc, C57BL/6 mice were subjected to electric foot shock, and the NAc was collected using a biopsy punch after the stimuli. The phosphorylation of ARHGEF2 at T103 and of ARHGAP21 at S875 was immediately higher in foot-shocked mice than in non-foot-shocked control mice (Fig. 3e). The Rho-kinase substrate MYPT1 was also phosphorylated at T694 and T852 in the NAc after the electric shock (Fig. 3f). Whereas the level of phosphorylation was reduced by the intraperitoneal injection of fasudil (a Rho-kinase inhibitor that passes through the blood–brain barrier) (Fig. 3f). To determine the cellular population of phosphorylated MYPT1 evoked by electric foot shock, we monitored the immunofluorescence of phospho-MYPT1 (T694) in the NAc of Drd1-tdTomato/Drd2-YFP double-transgenic mice, in which D1R-MSNs expressed tdTomato and D2R-MSNs expressed YFP. Electric foot shock significantly increased the number of cells positively labelled with phospho-MYPT1 (T694) in both D2R-MSNs and D1R-MSNs (Fig. 3g). Electric shock-induced phosphorylation of MYPT1 was inhibited by fasudil in D2R-MSNs and D1R-MSNs (Fig. 3g). The inhibitory effect of fasudil was weaker in D1R-MSNs than in D2R-MSNs. This may be due to phosphorylation of MYPT1 by other kinases (e.g. MRCK) in addition to Rho-kinase in D1R-MSN. These results suggest that aversive stimuli activate the CaMKII–RhoA–Rho-kinase pathway in D2R-MSNs of the NAc.

### Rho-kinase is involved in aversive learning and synaptic plasticity in D2R-MSNs

Several pharmacological studies have shown that Rho-kinase inhibition impairs memory formation, including long-term spatial memory in the hippocampus^35^, and fear conditioning memory in the lateral amygdala^36^. However, whether Rho-kinase regulates aversive learning remains unclear. To examine whether Rho-kinase regulates aversive learning, a passive avoidance test, a fear-motivated test classically used to assess memory in laboratory animals, was performed. Rho-kinase is composed of two subfamilies, ROCK1 and ROCK2, both of which show similar catalytic activity^37^. Heterozygous *Rock1* and *Rock2* double loxP-flanked (floxed) mice (*Rock1*^fl/+^;*Rock2*^fl/+^) were generated by crossing homozygous *Rock1* floxed mice (*Rock1*^fl/fl^)^38^ with homozygous *Rock2* floxed mice (*Rock2*^fl/fl^) (Supplementary Fig. 3a-e). AAV-CaMKII-Cre was injected into the NAc of homozygous *Rock1* and *Rock2* double floxed mice (*Rock1*^fl/fl^;*Rock2* ^fl/fl^) to locally deplete both ROCK1 and ROCK2. The expression of Cre in the NAc significantly decreased ROCK1 and ROCK2 protein expression in the NAc (Fig. 4a). Compared with control treatment, the localized knockout of both ROCK1 and ROCK2 in the NAc significantly reduced step-through latency (Fig. 4b). We further investigated whether the inhibition of Rho-kinase in D1R-MSNs or D2R-MSNs regulated aversive learning. The dominant negative (DN) form of ROCK2 (RB/PH(TT)) can inhibit the activity of ROCK1 and ROCK2. AAV-Flex-DN-ROCK2 was injected into the NAc of Drd1a-Cre or Adora2a-Cre transgenic mice^33, 34^ to specifically express DN-ROCK2 in the D1R-MSNs or D2R-MSNs. Step-through latency was significantly lower under DN-ROCK2 expression conditions than under control conditions among Adora2a-Cre mice (Fig. 4c) but not among Drd1a-Cre mice (Fig. 4c). These results indicate that Rho-kinase in D2R-MSNs is involved in aversive learning.

**Fig. 4:**
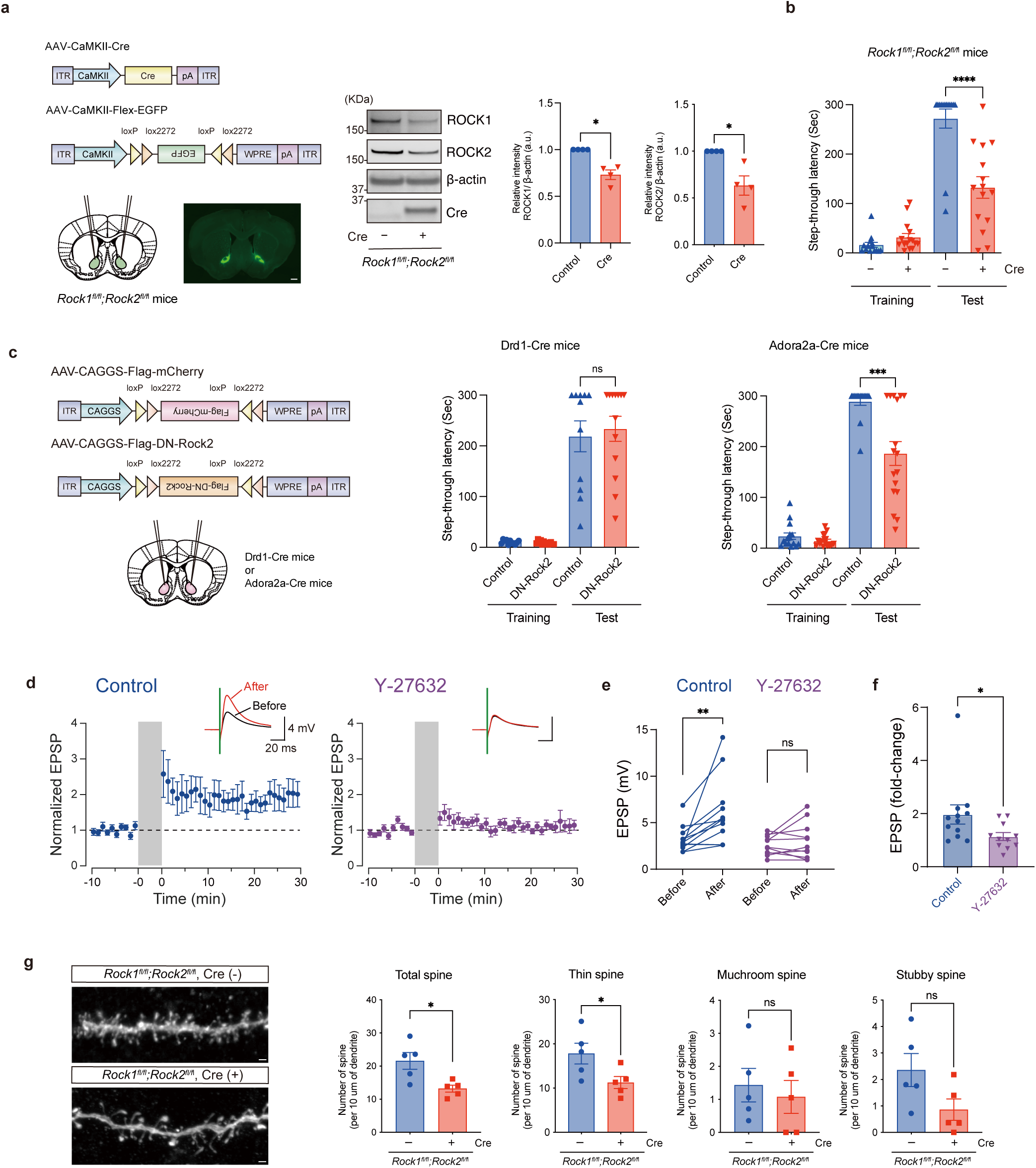
Rho-kinase is involved in aversive learning and synaptic plasticity in D2R-MSNs **(a)** AAV-CaMKII-Flex-EGFP and AAV-CaMKII-Cre were injected into the NAc of homozygous *Rock1* and *Rock2* double floxed mice (*Rock1^fl/fl^*;*Rock2 ^fl/fl^*). Representative coronal brain slices show the expression of EGFP 3 weeks after AAV injection (left panel). Scale bar=500 µm. AAV-mediated knockout of *Rock1 and Rock2* decreased the ROCK1 and ROCK2 protein levels in the NAc (right panel). Error bars indicate the mean ± SEM of four independent experiments. Mann Whitney test. **p<0.05. **(b)** Loss of ROCK1 and ROCK2 in the NAc suppressed aversive learning. Error bars indicate the mean ± SEM (Cre (-) n=15; Cre (+) n=14). Mann-Whitney U test. ****p<0.0001. **(c)** Inhibition of ROCK2 in D2R-MSNs suppressed aversive learning. AAV-CAGGS-Flex-mCherry and AAV-CAGGS-Flex-Flag-ROCK2-RB/PH(TT) (DN-ROCK2) were injected into the NAc of Drd1-Cre or Adora2a-Cre mice. The expression of DN-ROCK2 in D1R-MSNs did not alter step-through latency. Error bars indicate the mean ± SEM (Control, n=12; DN-ROCK2, n=13). The expression of DN-ROCK2 in D2R-MSNs reduced the step-through latency. Error bars indicate the mean ± SEM (Control, n=15; DN-ROCK2, n=13). Mann-Whitney U test. ***p<0.001. ns: not significant. **(d)** Averaged time course of excitatory postsynaptic potentials (EPSPs) recorded from D2R-MSNs in the NAc. An STDP protocol (grey shading) induced LTP in the control condition (n=12 cells, in the presence of 0.21% DMSO) but not in the presence of the Rho-kinase inhibitor Y-27632 (20 µM, n=11 cells). Insets show the grand average traces at the baseline (before) and 25–30 min after the STDP protocol application (After). A green bar in the inset indicates stimulation. **(e)** Quantification of mean EPSC amplitudes. Wilcoxon signed rank test (Control) or paired t test (Y-27632). **p<0.01. ns: not significant. **(f)** Quantification of fold-changes of EPSP amplitude induced by the STDP protocol application. Mann–Whitney U test. *p<0.05. **(g)** Loss of ROCK1 and ROCK2 in the NAc reduced spine density. The left panels show representative confocal images of viral-mediated EGFP expression in the secondary dendrites of D2R-MSNs. Scale bar=1 µm. The right panels show the quantification of the spine number (Total spine, Shin spine, Mushroom spine, and Stubby spine). Data present the mean ± SEM (n=5 cells per each condition, total 485 spines). Unpaired t test. *p<0.05. ns: not significant.

The long-term plasticity of synapses onto accumbal D2R-MSNs is likely involved in forming aversive memory. We therefore investigated whether Rho-kinase plays a role in long-term synaptic plasticity in the NAc. Application of a spike-timing-dependent plasticity (STDP) protocol^39^ induced long-term potentiation (LTP) at excitatory synapses onto D2R-MSNs in the NAc (Fig. 4d, e). Treatment of slices with a Rho-kinase inhibitor (Y-27632) significantly attenuated LTP at the late phase (25–30 min after the cessation of STDP protocol, p=0.0268) (Fig. 4d-f). The effect of Y-27632 on the early phase of LTP at 0–5 min after STDP induction was not significant (p=0.1507). These results are consistent with previous reports that Rho-kinase inhibitor prevented hippocampal LTP stabilization^40^. Considering that actin assembly inhibitors destabilize LTP without affecting the initial expression of LTP^41^, our results suggest that Rho-kinase is necessary for maintaining potentiated synapses through stabilization of the actin cytoskeleton of the dendritic spines^42^. To confirm this, the effect of Rho-kinase activity on structural plasticity of spine was investigated using homozygous *Rock1* and *Rock2* double floxed mice (*Rock1^fl/fl^*;*Rock2 ^fl/fl^*). The analysis of dendritic spines revealed that the disruption of both ROCK1 and ROCK2 reduced the total spine density (Fig. 4g), suggesting that Rho-kinase regulated the structural plasticity of the dendritic spine in D2R-MSNs in the NAc.

### Identification of SHANK3 as a novel Rho-kinase substrate

Based on our findings, we next tried to identify the substrate phosphorylated by Rho-kinase in the regulation of synaptic plasticity and aversive learning. To explore the putative substrates for Rho-kinase, phosphoproteomic analysis was performed using the KIOSS method. A phosphatase inhibitor was used to increase the phosphorylation of many proteins, and a specific kinase inhibitor was used to suppress targeted kinase-induced phosphorylation enhanced by a phosphatase inhibitor^14^. When striatal/accumbal slices were treated with calyculin-A (CLA, a specific inhibitor of type 1 and 2A phosphatases) and Y27632 (an *in vitro* Rho-kinase inhibitor), CLA induced the phosphorylation of MYPT1, whereas Y-27632 inhibited the CLA-induced phosphorylation of MYPT1 (Supplementary Fig. 4a). The ζ isotype of 14-3-3 is known to be involved in synaptic morphology^43^. Therefore, to explore the synaptic phosphorylated proteins, striatal/accumbal lysates treated with CLA and/or Y-27632 were added to affinity beads coated with GST-14-3-3 ζ. The bound proteins were digested with trypsin and analyzed with LC-MS/MS (Supplementary Fig. 4b). The identified proteins were considered candidate substrates for Rho-kinase if they fulfilled three criteria: (1) the phosphorylation level of the protein based on the ion intensity was increased more than 2-fold by treatment with CLA (2) the phosphorylation of the protein was decreased by treatment with Y-27632, and (3) The phosphorylation motif is consistent with the consensus for Rho-kinase [(R/K)XX(pS/pT)] or [(R/K)X(pS/pT)]. A total of 221 proteins were identified as candidate substrates for Rho-kinase (Supplementary Fig. 4c and Supplementary Table 3). Pathway analysis using the Reactome database revealed that ‘protein–protein interactions at synapses’ is one of the Rho-kinase-related pathways (Supplementary Fig. 4d). Synaptic structural plasticity is regulated by protein–protein interactions and post-translational modifications of postsynaptic density (PSD) scaffolding proteins^44^. Among the identified candidate substrates of Rho-kinase, we focused on SH3 and multiple ankyrin repeat domains 3 (SHANK3). This is because SHANK3 is a scaffolding protein in the PSD of excitatory synapses that interconnects NMDARs and AMPARs via complexes with discs large homolog-associated protein 3 (DLGAP3) and postsynaptic density protein 95 (PSD95)^45^. Moreover, SHANK3 is thought to interact with actin-binding proteins, including cortactin, to promote actin polymerization in spines and to link actin filaments to NMDARs and AMPARs^45, 46^. In addition, SHANK3 knockdown or knockout reduces the density, maturation, and actin levels of dendritic spines^47, 48, 49^.

The mass spectrometry results identified three sites near the SH3-PDZ domain (T551, S694, and S781) as candidate sites for phosphorylation by Rho-kinase (Fig. 5a). To confirm whether these SHANK3 phosphorylation sites were phosphorylated by Rho-kinase, we performed an in vitro kinase assay. Rho-kinase-CAT phosphorylated the SHANK3-SH3-PDZ domain *in vitro*, whereas the level of phosphorylation was approximately 60% lower in SHANK3-SH3-PDZ-3A (in which the T551, S694, and S781 sites were mutated to alanine) (Fig. 5b). These results indicate that Rho-kinase directly phosphorylates SHANK3 at T551, S694, and S781 *in vitro*. We then produced phosphorylation-specific antibodies against phosphorylated T551, S694, and S781. These antibodies recognized SHANK3, which was phosphorylated by Rho-kinase, but not the non-phosphorylated SHANK3 (Supplementary Fig. 2c-e). The treatment of striatal/accumbal slices with high K^+^ or NMDA induced the phosphorylation of SHANK3 at T551, S694, and S781, and this was inhibited by pretreatment with a Rho-kinase inhibitor (Y-27632) (Fig. 5c). These results indicate that NMDA-induced Rho-kinase activation mediates the phosphorylation of SHANK3 in striatal/accumbal slices. We next examined whether electric foot shock induced the phosphorylation of SHANK3 in the NAc. The phosphorylation of SHANK3 at T551, S694, and S781 was significantly elevated after electric shock and was reduced by the administration of fasudil (Fig. 5d). These results indicate that electric shock induces the Rho-kinase-mediated phosphorylation of SHANK3 in the NAc.

**Fig. 5:**
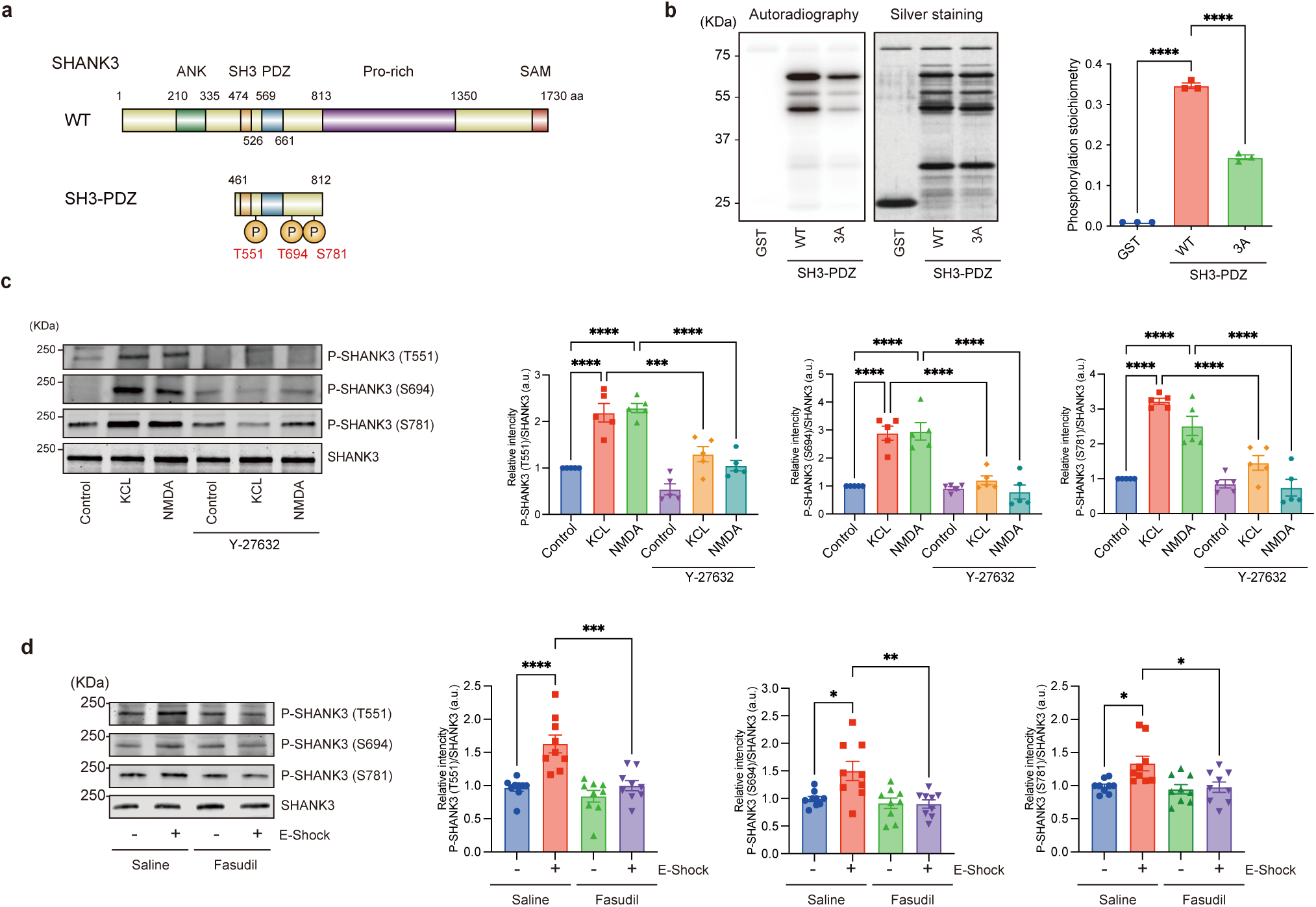
Identification of SHANK3 as a novel Rho-kinase substrate **(a)** Schematic of SHANK3 domain structure and phosphorylation sites. **(b)** Rho-kinase phosphorylated SHANK3-SH3-PDZ *in vitro*. Data represent the mean ± SEM of three independent experiments. One-way ANOVA followed by Tukey’s multiple comparisons test. ****p<0.0001. **(c)** Rho-kinase phosphorylated SHANK3 at T551, S694, and S781 downstream of NMDARs in the striatum/NAc. Striatal/accumbal slices were pretreated with Y-27632 (20 µM) for 1 h, treated with KCL (40 mM) or NMDA (100 µM) for 15 s, and subjected to immunoblot analysis. Data present the mean ± SEM of five independent experiments. Two-way ANOVA followed by Tukey’s multiple comparisons test. ***p<0.001, ****p<0.0001. **(d)** Electric foot shock induced the phosphorylation of SHANK3 at T551, S694, and S781 in vivo. C57BL/6J mice were intraperitoneally injected with saline or fasudil (20 mg/kg) for 15 min and then subjected to electric foot shock (0.7 mA, 3 s). Data present the mean ± SEM (n=9 mice per each condition). Two-way ANOVA followed by Tukey’s multiple comparisons test. *p<0.05, **p<0.01, ***p<0.001, ****p<0.0001.

### SHANK3 phosphorylation is important for spine formation and aversive learning

To examine whether SHANK3 was involved in aversive learning, a passive avoidance test was performed using homozygous *Shank3* floxed mice (*Shank3*^fl/fl^) (Supplementary Fig. 5a-c). AAV-CaMKII-Cre was injected into the NAc of homozygous *Shank3* floxed mice (*Shank3*^fl/fl^) to locally deplete SHANK3. The expression of Cre in the NAc significantly decreased SHANK3 protein expression in the NAc (Fig. 6a). Step-through latency was significantly lower under localized knockout of SHANK3 in the NAc than under control conditions (Fig. 6b), indicating that SHANK3 in the NAc was involved in aversive learning.

**Fig. 6:**
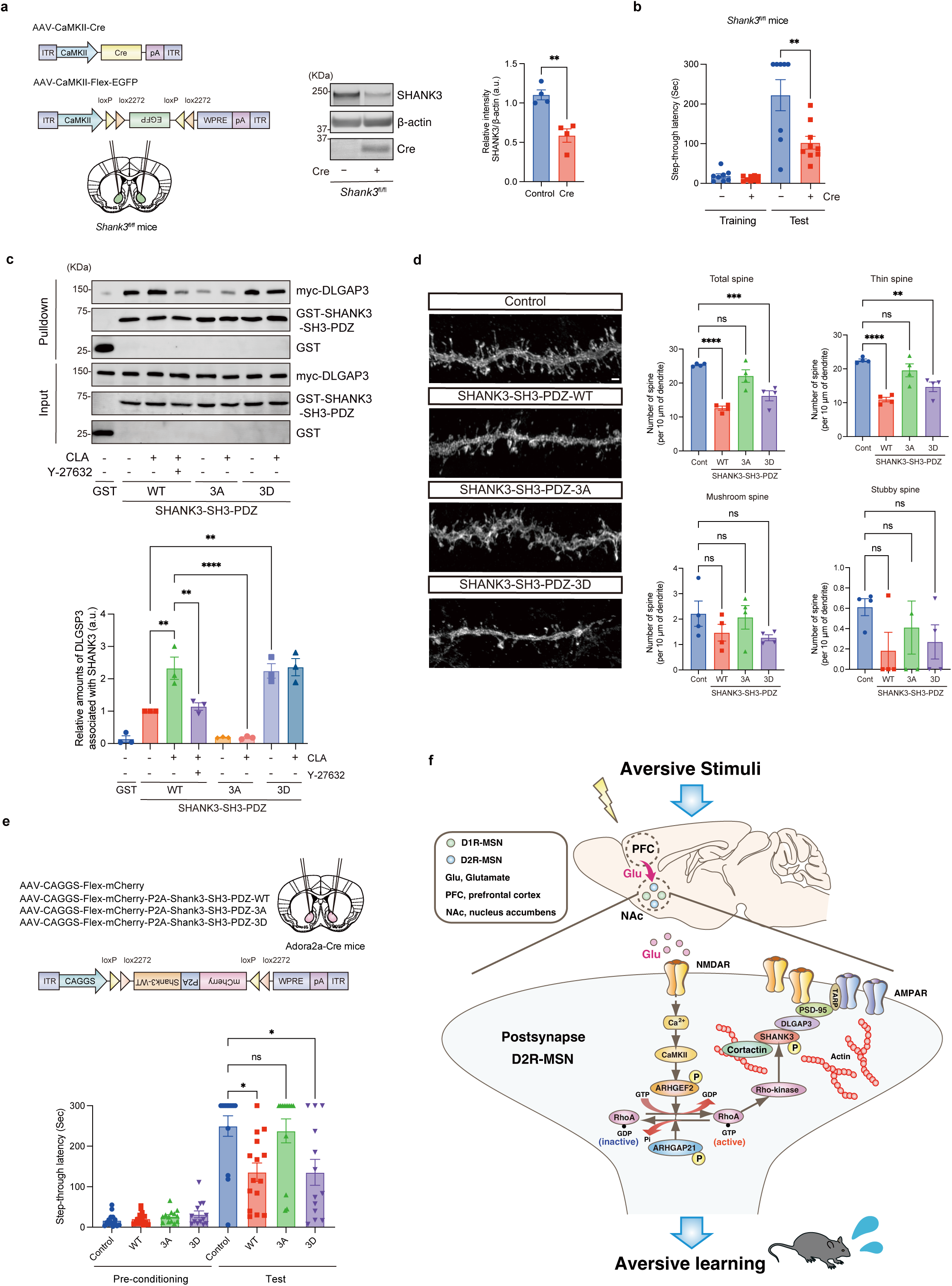
SHANK3 phosphorylation is important for spine growth and aversive learning. **(a)** AAV-mediated knockout of *Shank3* decreased the SHANK3 protein levels in the NAc. Error bars indicate the mean ± SEM of four independent experiments. Unpaired t test. **p<0.01. **(b)** Loss of SHANK3 in the NAc suppressed aversive learning. Error bars indicate the mean ± SEM (Cre (-) n=8; Cre (+) n=9). Mann-Whitney U test. **p<0.01. **(c)** Rho-kinase-dependent interaction between SHANK3 and DLGAP3 in COS7 cells. COS7 cells were transfected with indicated plasmids and subsequently treated with calyculin-A (CLA, 350 nM) and/or Y-27632 (20 µM), followed by pulldown with glutathione 4B beads. Data present the mean ± SEM of three independent experiments. One-way ANOVA followed by Tukey’s multiple comparisons test. **p<0.01, ****p<0.0001. **(d)** The expression of DN-SHANK3 in D2R-MSNs reduced spine density. AAV-CaMKII-Cre and AAV-CAGGS-Flex-mCherry-P2A-SHANK3-SH3-PDZ-WT, −3A, or −3D were injected into the NAc of D2R-YFP mice. The left panels show representative confocal images of viral-mediated mCherry expression in the secondary dendrites of D2R-MSNs. Scale bar=1 µm. The right panels show the quantification of the spine number. Data present the mean ± SEM (n=4 cells per each condition, total 865 spines). One-way ANOVA followed by Tukey’s multiple comparisons test. **p<0.01, ***p<0.001, ****p<0.0001. ns: not significant. **(e)** The expression of DN-SHANK3 in D2R-MSNs suppressed aversive learning. Indicated AAV was injected into the NAc of Adora2a-cre mice. Mice were subjected to a passive avoidance test. Error bars indicate the mean ± SEM (Control, n=14; WT, n=15; 3A, n=13; 3D, n=13). Kruskal-Wallis test followed by Dunnett’s multiple comparisons test. *p<0.05. ns: not significant. **(f)** Proposed model of the downstream effects of the CaMKII-RhoA-Rho-kinase pathway. When mice are exposed to aversive stimuli, glutamate acts on NMDA receptors and activates the CaMKII-RhoA-Rho-kinase pathway via Ca^2+^ influx in D2R-MSNs. Rho-kinase phosphorylates SHANK3, promotes actin polymerization, and stabilizes receptors to the membrane, thereby regulating synaptic structural plasticity and promoting aversive learning.

The interaction between SHANK3 and NMDARs is mediated by the binding of SHANK3 to DLGAP3, which leads to the formation of the PSD95/DLGAP3/SHANK3 complex^45^. To examine whether the phosphorylation of SHANK3 modulated its interaction with DLGAP3, a GST pulldown assay was performed using phospho-deficient mutant of SHANK3 (SHANK3-SH3-PDZ-3A) and phospho-mimic mutant of SHANK3 (i.e. SHANK3-SH3-PDZ-3D, in which the T551, S694, and S781 sites were mutated to aspartic acid). SHANK3-SH3-PDZ-WT interacted with DLGAP3, and treatments with CLA increased the interactions between SHANK3-WT and DLGAP3, but this interaction was inhibited by pretreatment with a Rho-kinase inhibitor (Y-27632). SHANK3-SH3-PDZ-3A lost its ability to bind with DLGAP3, whereas SHANK3-SH3-PDZ-3D showed stronger binding with DLGAP3 compared with SHANK3-SH3-PDZ-WT (Fig. 6c). These results indicate that the phosphorylation of SHANK3 by Rho-kinase increases its interaction with DLGAP3.

It has been reported that truncated SHANK3 containing SH3-PDZ acts as a DN and reduces the density of dendritic spines^49^. Therefore, we next investigated whether the expression of SHANK3-SH3-PDZ affects the spine density in D2R-MSNs of the NAc. The analysis of dendritic spines revealed that the expression of SHANK3-SH3-PDZ-WT, which may inhibit the interaction between SHANK3 and DLGAP3 (Fig. 6c) in D2R-MSNs, reduced the total spine density (Fig. 6d). This suggested that SHANK3-SH3-PDZ-WT had a DN effect on spine formation in D2R-MSNs. SHANK3-SH3-PDZ-3D expression in D2R-MSNs reduced spine density to a level similar to that of SHANK3-SH3-PDZ-WT expression (Fig. 6d). In contrast, the expression of SHANK3-SH3-PDZ-3A, which may not inhibit the SHANK3–DLGAP3 interaction (Fig. 6c), did not reduce spine density (Fig. 6d). This suggests that the Rho-kinase-induced phosphorylation of SHANK3 in D2R-MSNs is important for spine formation.

Finally, we investigated whether the expression of the SHANK3 mutant in D2R-MSNs affected aversive learning. AAV-Flex-mCherry-P2A-SHANK3-SH3-PDZ mutants were injected into the NAc of Adora2a-Cre mice. The expression of SHANK3-SH3-PDZ-WT in D2R-MSNs significantly attenuated step-through latency (Fig. 6e), indicating that SHANK3 dysfunction in D2R-MSNs affected aversive learning and memory. SHANK3-SH3-PDZ-3D expression in D2R-MSNs significantly decreased step-through latency, whereas SHANK3-SH3-PDZ-3A expression did not reduce step-through latency (Fig. 6e). Collectively, these results suggest that the phosphorylation of SHANK3 in D2R-MSNs is important for aversive learning and memory.

## Discussion

In this study, we used our originally developed phosphoproteomic analysis to explore the phosphorylation signals downstream of NMDARs in the striatum/NAc. We identified 194 proteins, including ARHGEF2 and ARHGAP21, whose phosphorylation was stimulated by NMDA in the striatal/accumbal slices. CaMKII phosphorylated ARHGEF2 and stimulated its RhoGEF activity, thereby activating RhoA-Rho-kinase. Moreover, we identified 221 candidate substrates of Rho-kinase, including SHANK3 and demonstrated that NMDA activated the CaMKII-RhoA-Rho-kinase pathway to induce SHANK3 phosphorylation, thereby regulating dendritic spine formation and aversive learning (Fig. 6f).

### Phosphoproteomic analysis of NMDAR signaling

Our phosphoproteomic analysis identified RhoA-specific regulators, including ARHGEF2 and ARHGAP21, as candidate substrates for CaMKII downstream of NMDAR (Fig. 1h, i). Besides RhoA, activation of Rac and Cdc42 downstream of NMDAR is also known to be important for synaptic plasticity and learning. Our phosphoproteomic analysis also identified, several Rac/Cdc42 regulators, including Fgd3, Tiam1, Arhgap25, Arhgap32, Bcr, Git1, and Srgap3, as candidate substrates of CaMKII downstream of NMDAR (Fig. 1h). Recent studies have shown that Tiam1 is phosphorylated and activated by CaMKII^50^. Furthermore, knock-in mice with mutations in the CaMKII binding site of Tiam1 show impaired learning^50^. Taken together, these findings suggest that the cooperative regulation of Rac and Cdc42, as well as RhoA, is important for synaptic plasticity and learning. In addition to Rho GTPase regulators, we identified many candidate substrates for CaMKII, including postsynaptic proteins (Dlg2, Dlg4, Dlgap2, Dlgap3, Grin1, Grin2b, Grm5, Iqsec1, Iqsec2, Shank2, Shank3, and Syngap1 etc.), presynaptic proteins (Bsn, Pclo, and Rims1 etc.), protein kinases (Aak1, Braf, Brsk2, Camk2a, Camk2b, Camk2g, Cdk17, Mark1, Mark3, Mast3, Mink1, Prkcb, Speg, and Wnk2 etc.), and ion channels (Ano1, Cacna1b, Cacna1e, Cacnb1, Hcn2, Kcnh2, and Kcnq2 etc.) (Fig. 1h, Table 1 and Supplementary Table 1). Further studies of these candidate substrates may reveal novel mechanisms involved not only in the striatum/NAc, but also in the function of other brain regions.

Based on our recent phosphoproteomic analysis to explore PKC substrates, we comprehensively identified 116 proteins as candidate substrates for PKC^51^. Of these, 72 proteins including Rac regulators (e.g. Arhgef7, Bcr and Tiam1) were consistent with those identified by phosphoproteomics using NMDA and high K^+^ (Fig. 1h). PKC activates Rac via β-PIX phosphorylation and regulates synaptic plasticity and aversive learning via the PAK-LIMK-cofilin pathway downstream of acetylcholine^51^. Given that PKC is also activated by NMDAR stimulation and is implicated in synaptic plasticity^52^, some candidate substrates of PKC may also be involved in synaptic plasticity and memory formation.

### Role of CaMKII-RhoA-Rho-kinase pathway for aversive learning

This study established that Rho-kinase plays a critical role in the NAc during aversive learning. Aversive stimuli induced the Rho-kinase-mediated phosphorylation of MYPT1 in the NAc (Fig. 3f, g). Moreover, genetic inhibition of Rho-kinase in the NAc impaired aversive learning (Fig. 4b, c). Inhibition of Rho-kinase in D2R-MSN suppressed stabilization of LTP (Fig. 4d-f) and spine formation (Fig. 4g). Collectively, these results suggest that Rho-kinase in the D2R-MSN of NAc contributes to LTP stabilization through regulating actin polymerization in dendritic spine and plays an important role in aversive learning. Rho-kinase regulates cofilin activity through the phosphorylation of LIM kinase, thereby promoting actin polymerization in HeLa cells^53^. As the LIM-kinase-cofilin pathway plays an important role in the structural plasticity of dendritic spines^54^, it is conceivable that Rho-kinase regulates structural plasticity through the LIM-kinase-cofilin pathway. However, it is difficult to explain all the functions of Rho-kinase using only LIM-kinase. Therefore, to identify the substrates based on which Rho-kinase functions, we comprehensively explored the substrates of Rho-kinase in striatal/accumbal slices using the KIOSS method (Supplementary Fig. 4a, b). Based on our analysis, we identified a number of candidate substrates, including postsynaptic proteins (Dlg2, Dlgap2, Lrrc7, Shank3, Syngap, Synpo1, and Synpo3), presynaptic proteins (Bsn, Erc1, Erc2, Pclo, Syn1, and Syn3), G protein regulators (Agap2, Agfg1, Als2, Bcr, Fdg1, Nf1, Srgap1, Srgap2, Tiam1, Tiam2, and Tsc2), ion channels (Cacna1b, Cacna1e, Kcnb1, Kcnh7, Kcnq2, Hcn4, and Nav1), adaptor proteins (Amer2, Nyap1, and Nyap1), membrane traffic components (Epn2, Iqsec1, Numb1, Rab11fip2, Rims1, Rims2, Reep1, Reep3, Shisa7, and Syt7), cytoskeletal proteins (Kif1a, Kif21b, Klc2, Klc3, Klc4, Map2, Map4, and Phactr1), and protein kinases (Aak1, Abl1, Abl2, Cdk14, Cdk16, Cdk17, Dclk1, Mark1, Mark3, Mast3, Mink1, and Sik3) (Supplementary Fig. 4c and Supplementary Table 2). Further studies are required to examine whether these candidates are directly phosphorylated by Rho-kinase and to identify the types of functional changes caused by their phosphorylation.

SHANK3 is thought to promote spine formation by stimulating actin polymerization at the spine head^49^. However, the mechanism of the regulation of SHANK3 in the PSD remains unclear^45, 55^. Here, we found that Rho-kinase phosphorylated SHANK3 in SH3-PDZ domain (Fig. 5b) and consequently promoted its interaction with DLGAP3 (Fig. 6c). The expression of the DN of SHANK3 (SHANK3-SH3-PDZ) and its phosphomimetic form (SHANK3-SH3-PDZ-3D) in D2R-MSNs inhibited spine formation and aversive learning (Fig. 6d, e), whereas the expression of the non-phosphorylatable form (SHANK3-SH3-PDZ-3A) did not show such inhibitory effects (Fig. 6d, e). These results suggest that the Rho-kinase-mediated phosphorylation of SHANK3 promotes its association with NMDARs and AMPARs via DLGAP3 and connects these complexes with actin filaments for promoting spine formation and aversive learning.

Based on our findings and previous observations showing that glutamatergic inputs from the frontal cortex to the NAc play a pivotal role in aversive learning^31^, we propose a model of the CaMKII-RhoA-Rho-kinase pathway (Fig. 6f). When aversive stimuli are provided to mice, glutamate is released from the frontal cortex to the NAc at the nerve endings of the projected neurons. The NMDARs in D2R-MSNs are stimulated, and they promote Ca^2+^ inflow into cells, thereby increasing the intracellular Ca^2+^ concentration. The CaMKII-RhoA-Rho-kinase pathway is activated, and various substrates, including SHANK3, are phosphorylated by Rho-kinase in D2R-MSNs. As such, Rho-kinase controls the structural plasticity of dendritic spines and promotes aversive learning.

### Strengths and limitations of this study

Our study demonstrates that learning and memory is fundamentally a metabolic cascade, and our high-throughput phosphoproteomic methods provide a tool to unlock major conceptual advances in various biological signalling cascades hitherto impossible in conventional studies. Our methods open up the possibility of understanding the entire phosphorylation network in our brains and allows us to study more targets to get the whole picture.

Our findings provide novel insights into the mechanisms through which the substrates downstream of NMDA receptors regulate aversive learning in D2R-MSN of the striatum/NAc. However, in addition to D2R-MSN, D1R-MSN is also present in the striatum/NAc, and D1R-MSN also receives glutamatergic input from various brain regions to regulate reward behaviour. Whether the CaMKII-RhoA-Rho-kinase pathway regulates reward behaviour and learning in the D1R-MSN is still unclear and remains an interesting future question. Furthermore, glutamate-mediated neurotransmission exists in a variety of regions, including the hippocampus and amygdala. The function of the CaMKII-RhoA-Rho-kinase pathway in these other brain regions has not been elucidated. The phosphoproteomic analysis and information presented in this study, along with analytical tools such as phospho-specific antibodies, will allow us to elucidate phosphorylation signaling in multiple brain regions. Ultimately, this will be useful for the development of antipsychotic and anti-dementia drugs.

In conclusion, the combination of our large-scale phosphoproteomics methods and our KANPHOS database has allowed us to understand the signal transduction of NMDARs. Further studies are needed to understand the entire phosphorylation network in the brain.

## Methods

### EXPERIMENTAL MODEL AND SUBJECT DETAILS

#### Resources table

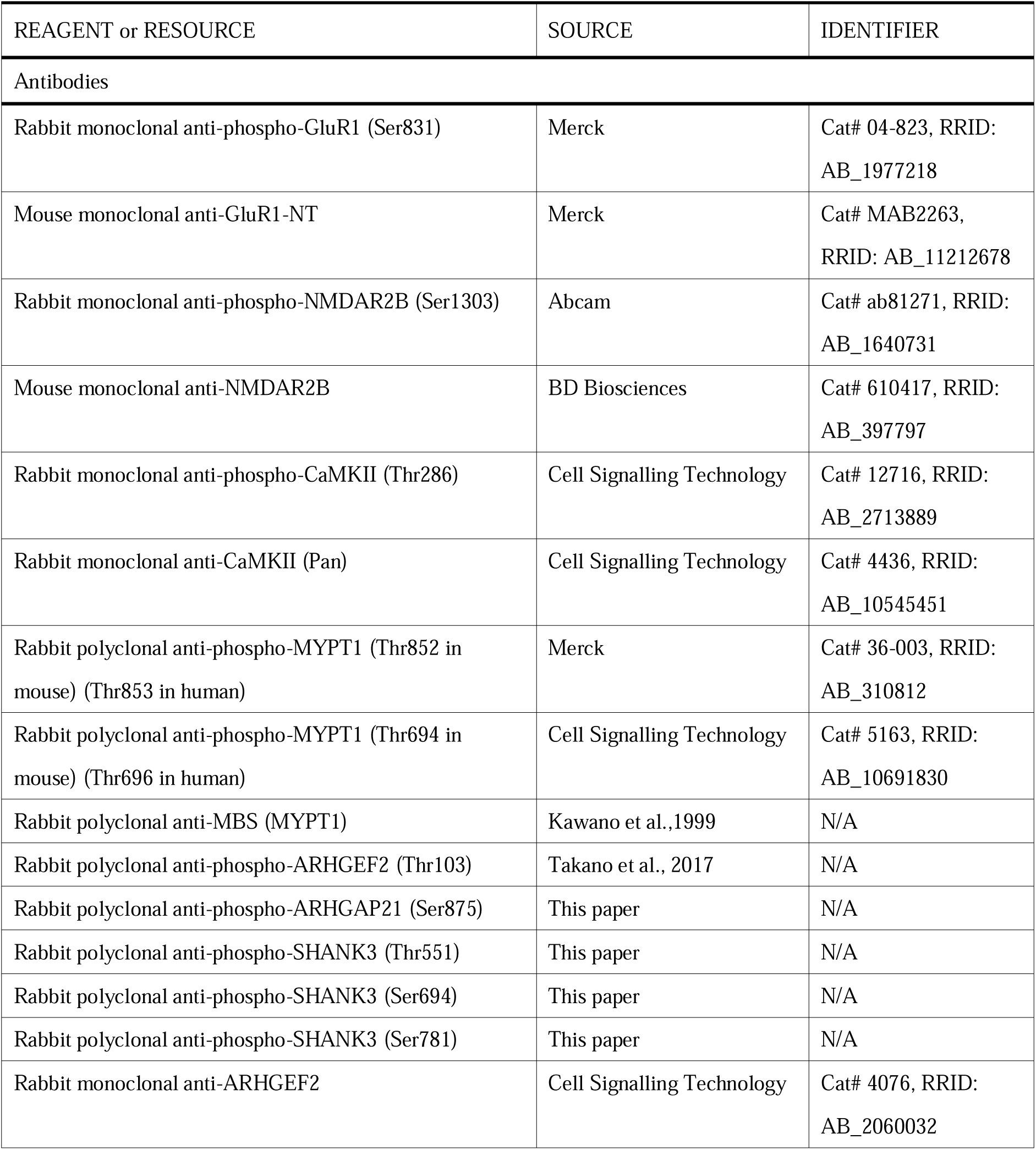
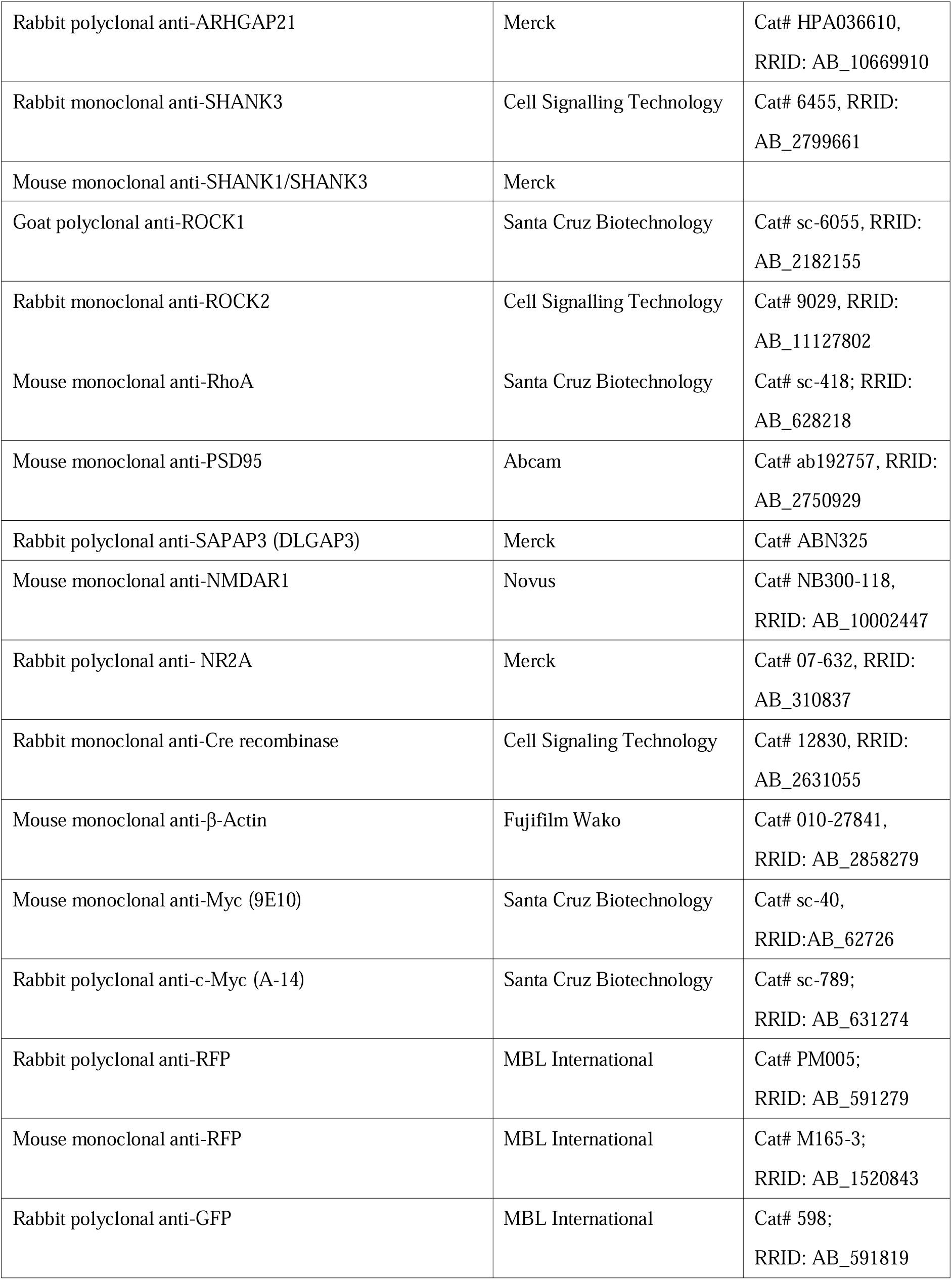
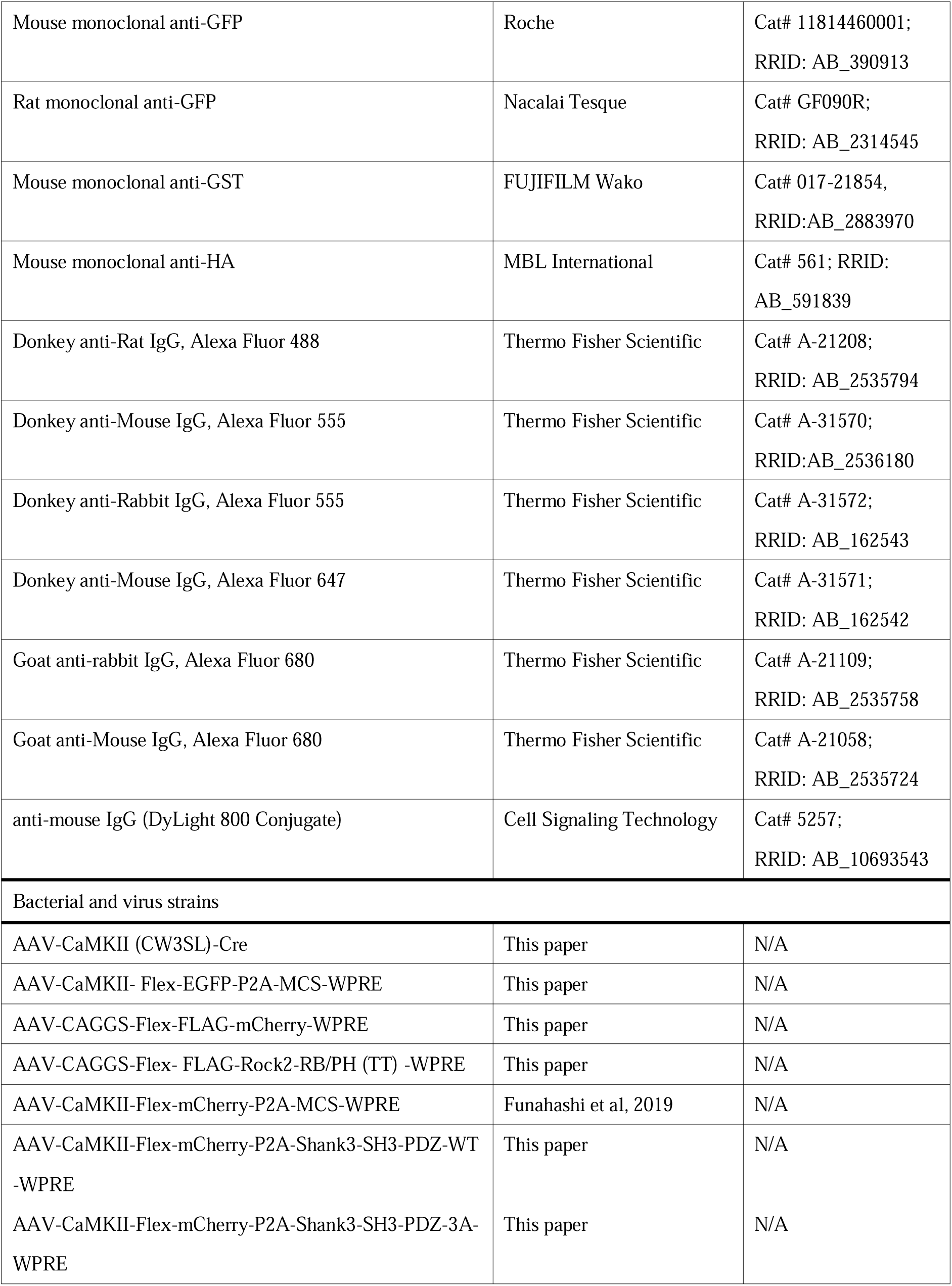
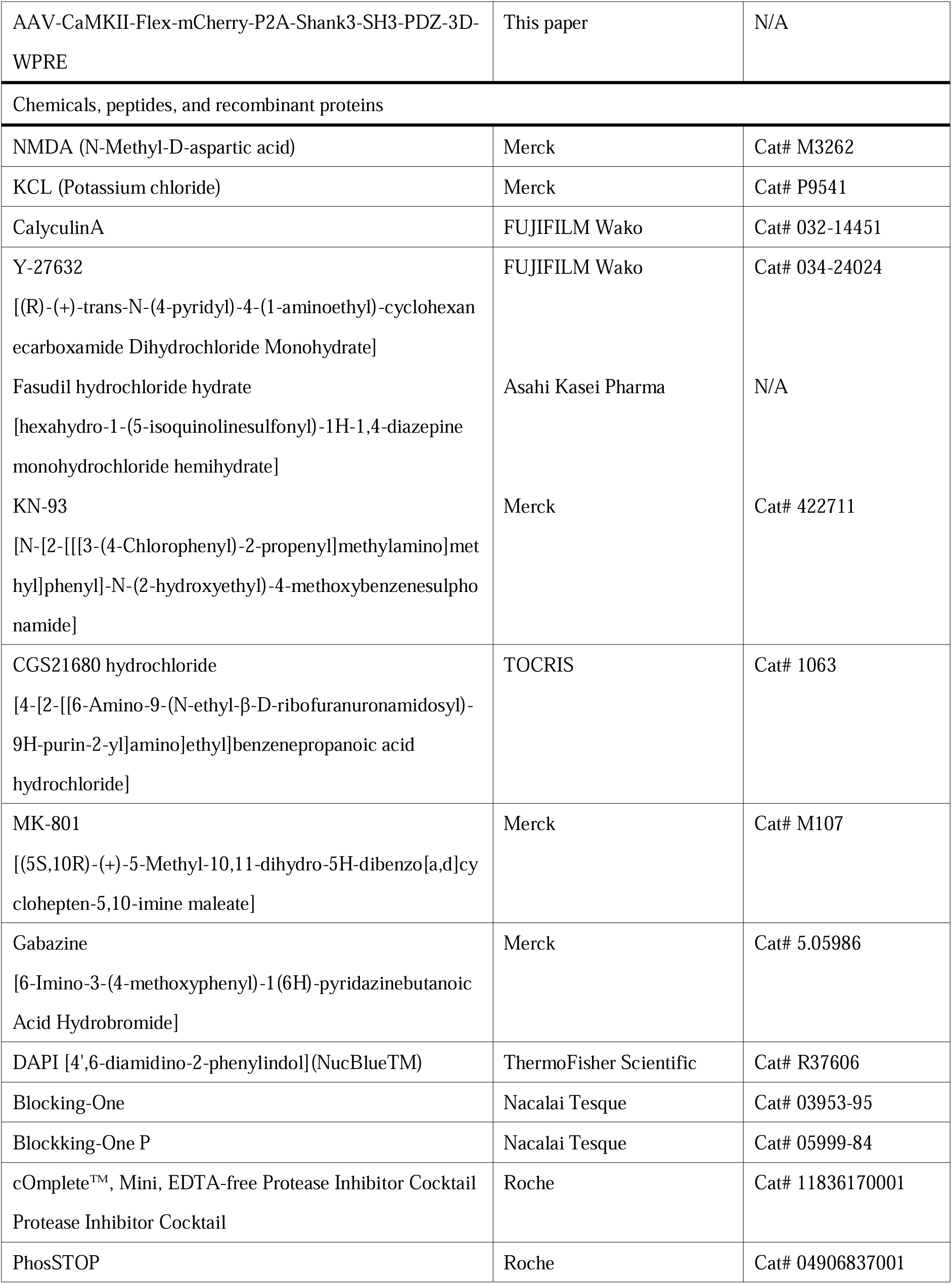
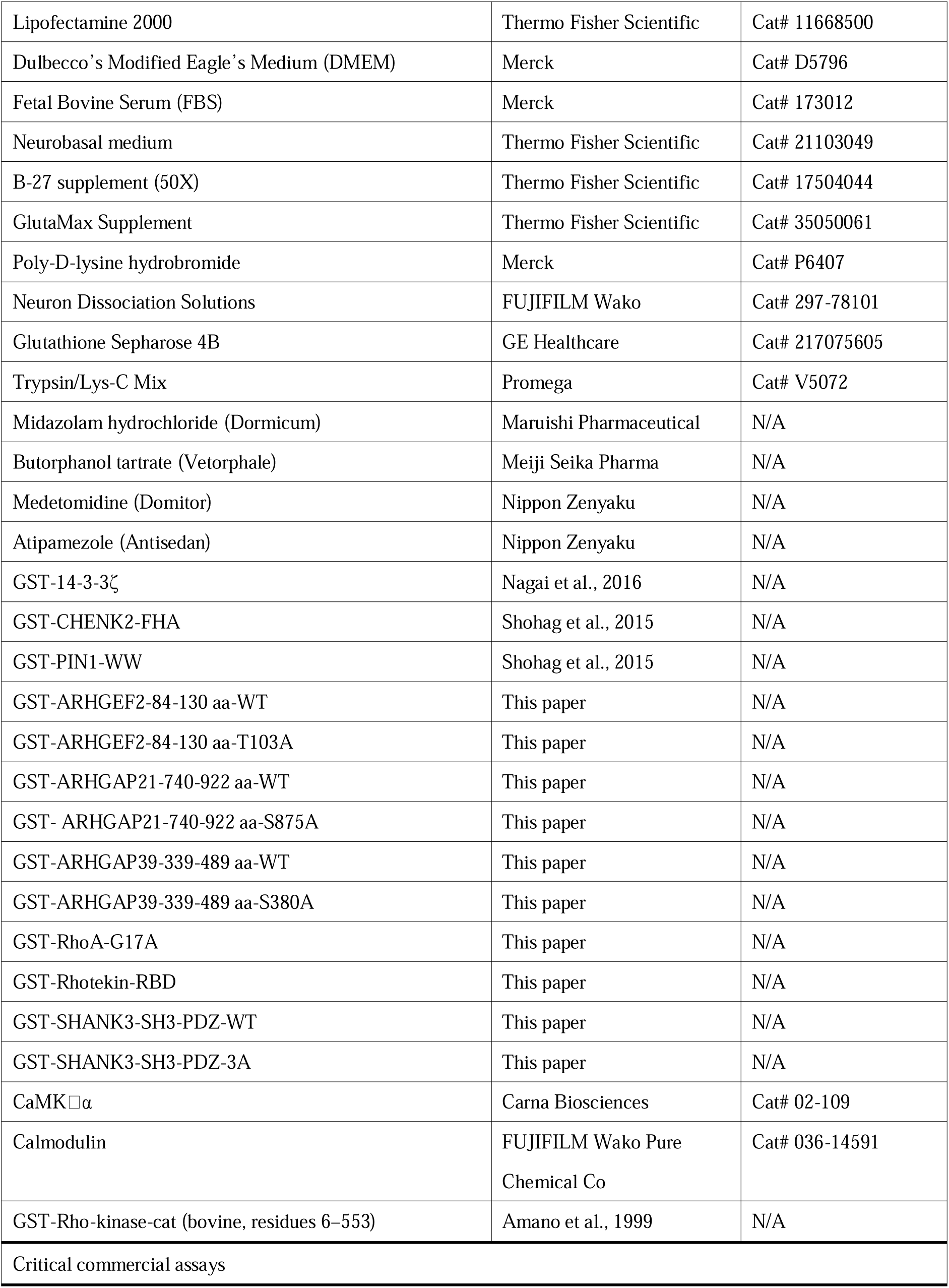
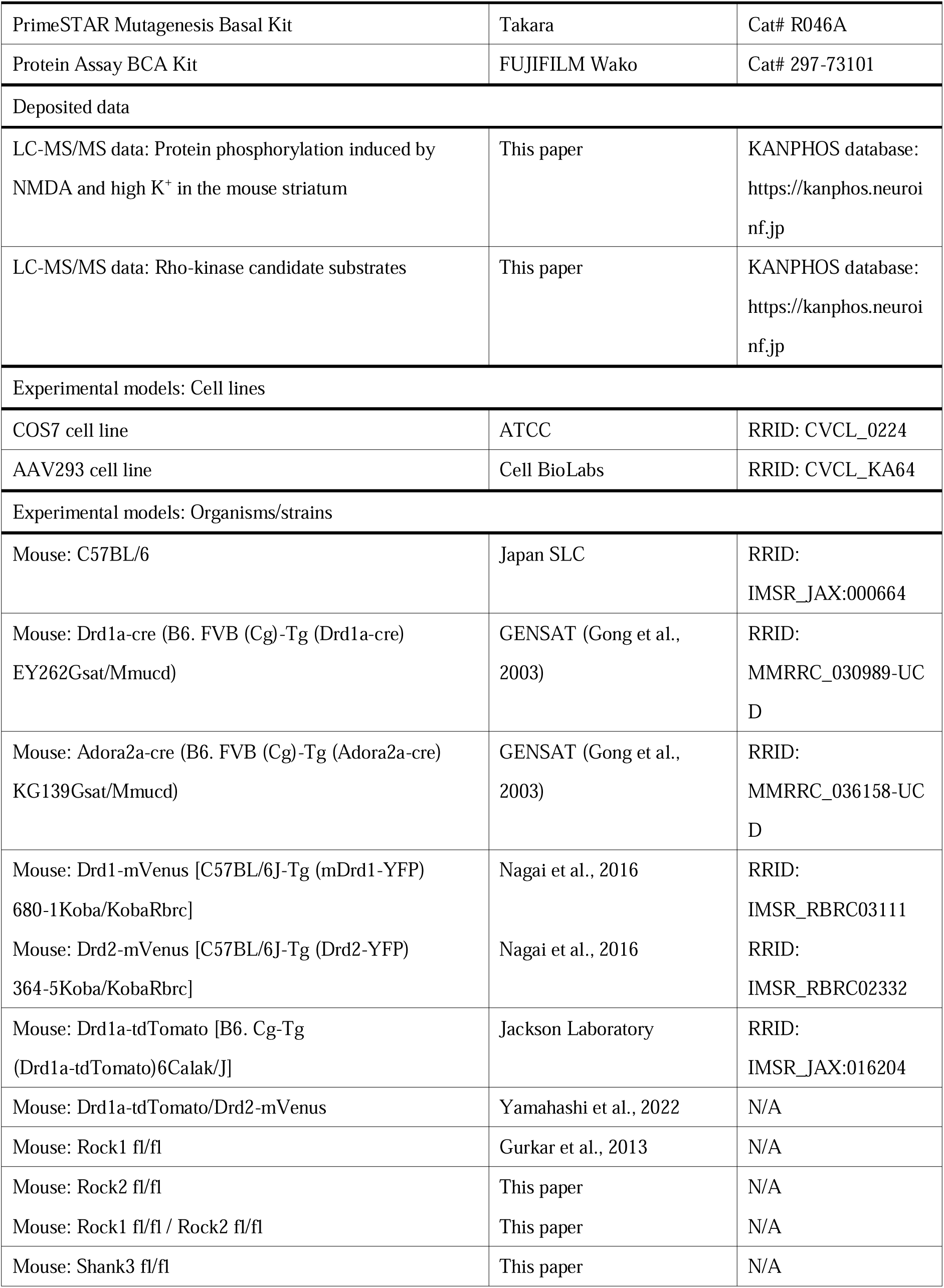
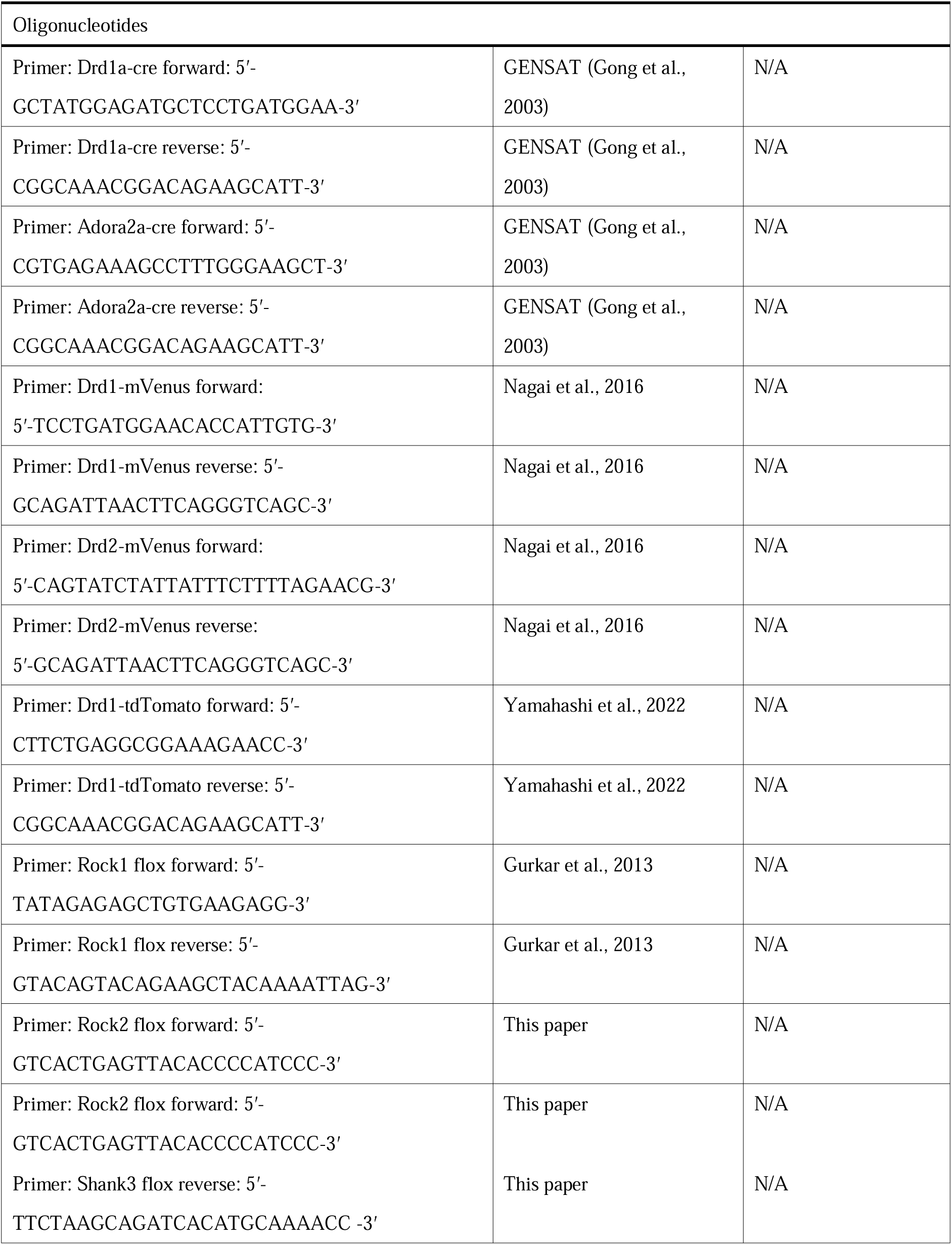
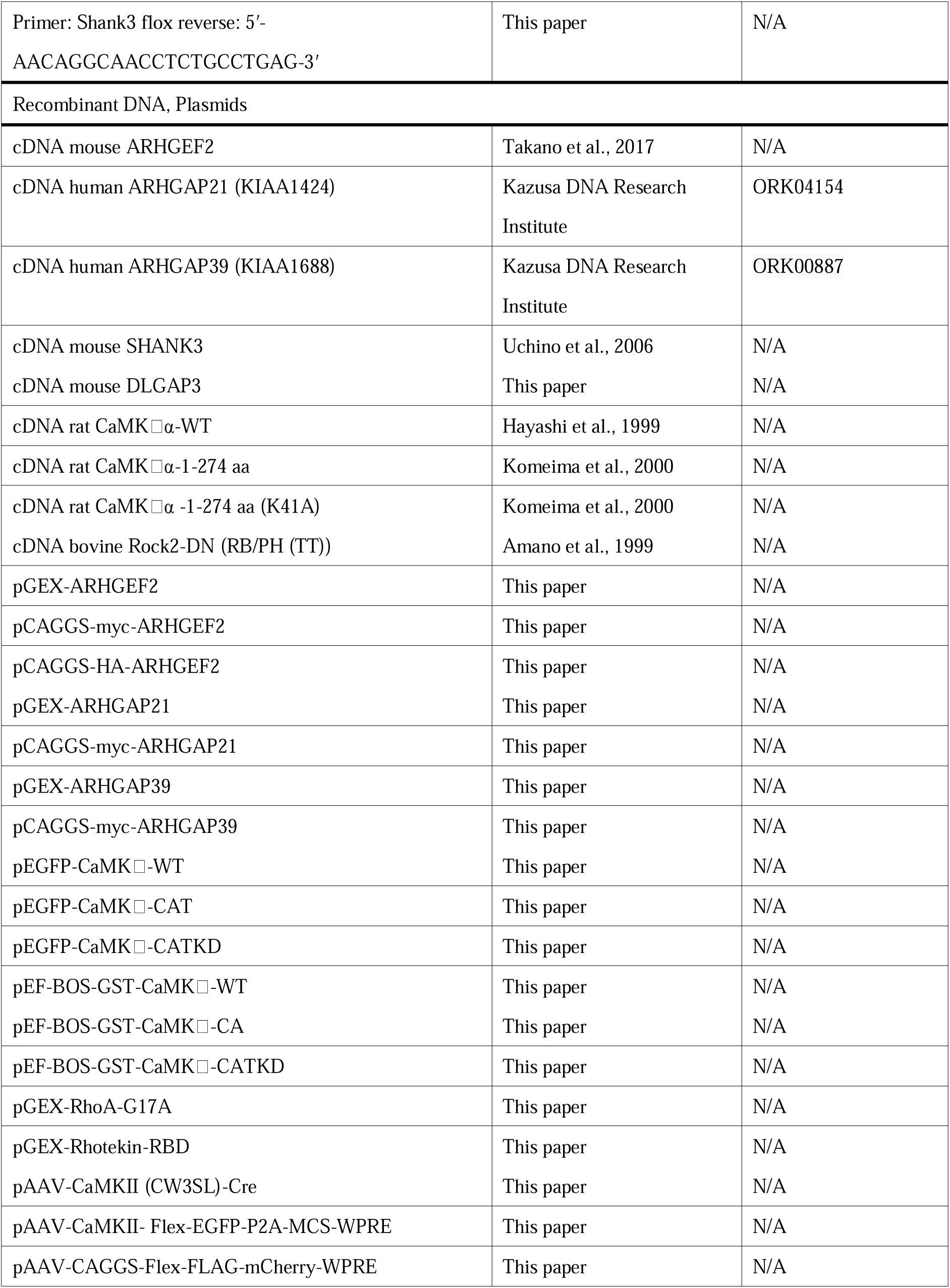
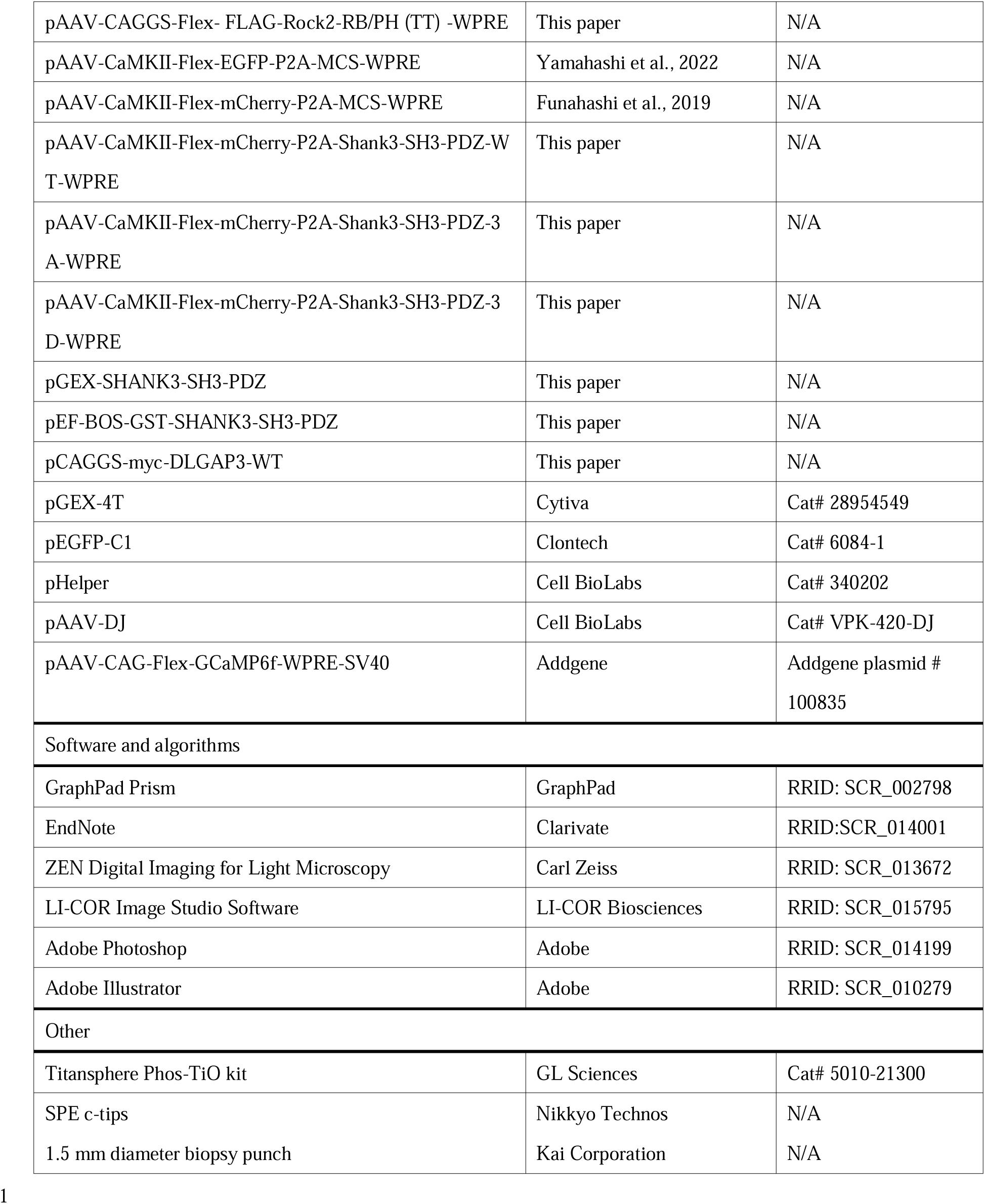

#### Mice

C57BL/6 (RRID: IMSR_JAX:000664) and ICR (RRID: IMSR_JAX:009122) mice were purchased from the Japan SLC (Shizuoka, Japan). *Rock1* floxed mice were provided by Dr. Young-Bum Kim (Harvard Medical School, Boston, MA, USA), as described previously^38^. *Rock2* floxed mice were generated as follows: C57BL/6N-Rock2tm1a(KOMP)Wtsi mice were generated from embryonic stem (ES) cells purchased from the International Mouse Phenotyping Consortium at the University of California, Davis (UC Davis, Davis, CA, USA). ES cells (EPD0492_1_B02 clone) were injected by RIKEN BRC (Tsukuba, Japan). The C57BL/6N-Rock2tm1a(KOMP)Wtsi ES cells used in this research project were generated by the trans-NIH Knock-Out Mouse Project (KOMP) and obtained from the KOMP Repository (www.komp.org). NIH grants to Velocigene at Regeneron Inc. (U01HG004085) and the CSD Consortium (U01HG004080) funded the generation of gene-targeted ES cells for 8500 genes in the KOMP Program, which were archived and distributed by the KOMP Repository at UC Davis and CHORI (U42RR024244). *Rock1* floxed mice and *Rock2* floxed mice were backcrossed six or more times on a C57BL/6N background (CLEA Japan, Inc.). Heterozygous *Rock1* and *Rock2* double-floxed mice (*Rock1*^fl/+^;*Rock2*^fl/+^) were obtained by crossing homozygous *Rock1* floxed mice (*Rock1*^fl/fl^) with homozygous *Rock2* floxed mice (*Rock2*^fl/fl^).

Drd1-mVenus (C57BL/6J-Tg(mDrd1-YFP)680-1Koba/KobaRbrc, RRID: IMSR_RBRC03111) and Drd2-mVenus (C57BL/6J-Tg(Drd2-YFP)364-5Koba/KobaRbrc, RRID: IMSR_RBRC02332) transgenic mice were generated as described previously^13^. The Drd1a-tdTomato line (B6.Cg-Tg(Drd1a-tdTomato)6Calak/J, RRID: IMSR_JAX:016204) was provided by the Jackson Laboratory (Bar Harbor, ME, USA). Drd1-tdTomato/Drd2-mVenus double-transgenic mice were generated by crossing heterozygous Drd1a-tdTomato mice with homozygous Drd2-mVenus mice. Drd1a-Cre mice on a C57BL/6 background (B6.FVB(Cg)-Tg(Drd1a-cre)EY262Gsat/Mmucd, RRID: MMRRC_030989-UCD) and Adora2a-Cre mice on a C57BL/6 background (B6.FVB(Cg)-Tg(Adora2a-cre)KG139Gsat/Mmucd, RRID: MMRRC_036158-UCD) provided by the Mutant Mouse Research and Resource Center at UC Davis. Heterozygous Drd1a-Cre and Adora2a-Cre male mice were obtained by crossing heterozygous or homozygous Drd1a-Cre and Adora2a-Cre male mice with C57BL/6 female mice.

*Shank3* floxed mouse line was produced by homologous recombination using the ES cell line RENKA, which was developed from the C57BL/6N strain (Mishina and Sakimura 2007). The targeting vector was constructed in accordance with the mouse genomic DNA databases contained from exon 11 to exon 12 of *Shank3* gene. The neomycin phosphotransferase gene (Neo) and the gene encoding fragment A of diphtheria toxin (DT-A) were included for positive and negative selection, respectively. A DNA fragment containing a 34-bp loxP sequence and Pgk1 promoter-driven Neo flanked by a pair of flippase recombinase target (FRT) sequences was inserted upstream of exon 12. The other loxP site was introduced downstream of exon 11. Thus, Cre loxP deletion would delete exons 11 and 12, resulting in a nonsense mutation in the *Shank3* gene.

Introduction of the targeting vector into mouse ES cells, screening for homologous recombinants with southern blot analysis, and production of chimeric mice were carried. The resultant chimeric mice were mated with C57BL/6N mice, and offspring [Shank3+/lox(neo)] were further crossed with Actb-Flp mice to remove the neo cassette. The flp gene was bred out in the next generation. After confirming deletion of the neo cassette, heterozygous (*Shank3* flox/+) mice were mated to generate homozygous (*Shank3* flox/flox) mice.

All mice were housed under a standard 12-h light/dark cycle (light phase 9:00–21:00) at a constant temperature of 23±1°C and were provided with free access to food and water. The male mice were aged 7–12 weeks and weighed 22–29 g (unless otherwise indicated).

All animal experiments were performed in accordance with the guidelines for the care and use of laboratory animals established by the Animal Experiments Committees of the Nagoya University Graduate School of Medicine (reference number: M210568-002) and Fujita Health University (reference number: AP20037). All experiments were approved by the appropriate committees and were conducted in compliance with the ARRIVE guidelines.

#### Cell lines

COS7 (species: *Cercopithecus aethiops*; sex: male; RRID: CVCL_0224; ATCC, Manassas, VA, USA) and AAV293 (species: *Homo sapiens*; sex: female; RRID: CVCL_KA64; Cell BioLabs, San Diego, CA, USA) cell lines were cultured in Dulbecco’s Modified Eagle’s medium (Merck, Kenilworth, NJ, USA) containing 10% foetal bovine serum (Merck). The cells were grown in a humidified atmosphere containing 5% CO_2_ at 37°C.

#### Primary cell cultures

Primary cultured striatal neurons were prepared from postnatal (P1−3) *Rock1/2* fl/fl or *Shank3* fl/fl mice (Japan SLC). The striatal tissues were digested with neuron dissociation solutions (FUJIFILM Wako, Tokyo, Japan) according to the manufacturer’s instructions. Dissociated neurons were seeded on coverslips or dishes coated with poly-D-lysine (PDL; Merck) and cultured in Neurobasal medium (Thermo Fisher Scientific, MA, USA) containing 10% FBS (Merck). One hour after plating, the medium was changed to Neurobasal medium containing B-27 supplement (Thermo Fisher Scientific) and 1 mM GlutaMAX (Thermo Fisher Scientific). Neurons were grown in a humidified atmosphere of 5% CO2 at 37°C.

### METHOD DETAILS

#### Antibodies

Rabbit poly-clonal anti-MBS (MYPT1) antibody was generated by use of GST-rat MBS-N-terminal domain (1-707 aa) as previously described^56^. Rabbit polyclonal antibodies against ARHGEF2 phosphorylated at Thr-103 was produced against phosphopeptide C+LLRNNpTALQSVS (Merck) as previously described^57^. Rabbit polyclonal antibodies against ARHGAP21 phosphorylated at S875 was produced against phosphopeptides C+TERSKpSYDEGL (Merck). Rabbit polyclonal antibodies against SHANK3 phosphorylated at T551, S694, and S781 were produced against the phosphopeptides C+LFRHYpTVGSYD, C+TLRSKpSMTAEL, and C+IPRTKpSVGEDE (FUJIFILM Wako), respectively. We purified the phosphorylation site-specific antibody using an affinity column as follows. First, to prepare an affinity column with a phosphopeptide, we placed a peptide in a Sulfolink coupling gel (Thermo Fisher Scientific) and blocked non-specific binding using cysteine buffer (50 mM L-cysteine HCl, 50 mM Tris-HCl, and 5 mM ethylenediaminetetraacetic acid (EDTA); pH 8.5). Next, antiserum was applied to the column, and it was rotated for 24 h at 4°C. After washing with phosphate-buffered saline, the antibody was eluted with 100 mM glycine solution (pH 2.5), and the pH was adjusted to 7.5.

#### Plasmid constructs

cDNA mouse ARHGEF2 was obtained as previously described^57^. cDNA mouse SHANK3 was obtained as previously described^58^. cDNA of bovine DN ROCK2 (RB/PH(TT)) was obtained as previously described^59^. cDNA human ARHGAP21 (KIAA1424) was purchased from Kazusa DNA Research Institute (Chiba, Japan). cDNA rat CaMKIIα-WT^60^, 1-274 aa (CAT)^61^, and 1-274 aa (K41A, CATKD)^61^ were provided from Dr. Y Watanabe (Showa Pharmaceutical University, Tokyo, Japan). We subcloned the PCR-amplified cDNA fragments of ARHGEF2, ARHGAP21 SHANK3, and DLGAP3 into pCR-Blunt-TOPO vector or pENTR-D/TOPO vector (Thermo Fisher Scientific). After sequencing, these fragments were further subcloned into appropriate commercial or homemade destination vectors according to the manufacturer’s instructions. The mutants ARHGEF2-T103A, ARHGAP21-S875A, SHANK3-3A (T551A/S694A/S781A), and SHANK3-3D (T551D/S694D/S781D) were generated with a PrimeSTAR Mutagenesis Basal Kit (Takara, Shiga, Japan). pGEX-14-3-3ζ, pGEX-CHECK2-FHA, and pGEX-PIN1-WW were generated as previously described^13, 15^. GST-tagged proteins were produced in BL21 (DE3) or Rosetta (DE3) *Escherichia coli* cells and purified on glutathione-Sepharose 4B beads (Cytiva, Marlborough, MA, USA). pAAV-CaMKII-Flex-mCherry-P2A-MCS-WPRE and pAAV-CaMKII-Flex-EGFP-P2A-MCS-WPRE were generated as previously described^62^.

#### Preparation/incubation of striatal slices

Striatal slices were prepared from male C57BL/6 mice as described previously^13, 51^. Briefly, mouse brains were removed and stored in Mg^2+^ free Krebs-HCO_3_-buffer (124 mM NaCl, 4 mM KCl, 26 mM NaHCO_3_, 1.5 mM CaCl_2_, 1.25 mM KH_2_PO_4_, and 10 mM D-glucose; pH 7.4). Coronal brain slices (thickness, 350 μm) were prepared using a VT1200S vibratome (Leica Microsystems, Wetzlar, Germany). The whole striatum was isolated from the brain slices, and the striatal slices were incubated at 30°C in Krebs-HCO_3_-buffer (Mg^2+^ free) with 10 μg/mL adenosine deaminase (Roche, Basal, Switzerland) for 30 min under constant oxygenation with 95% O_2_/5% CO_2_. The buffer was replaced with fresh Krebs-HCO_3_^-^ buffer (Mg^2+^ free) and preincubated for 30 min. The striatal slices were pretreated with the indicated inhibitors for 60 min and then stimulated with the indicated activators. After stimulation, the slices were snap-frozen in liquid nitrogen and stored at −80°C. The slices were lysed in lysis buffer (50 mM Tris/HCl, 1 mM EDTA, 1% SDS, phosphatase inhibitor cocktail (Roche, Indianapolis, IN, USA), and protease inhibitor cocktail (Roche); pH 7.4) and sonicated for 20 s. The protein concentration of the lysates was determined using a BCA assay (FUJIFILM Wako).

#### In vivo *s*ample preparation

*In vivo* sample preparation for immunoblot analysis was performed as described previously^63^. The mice were decapitated immediately after electric foot shock. Each mouse head was immediately immersed in liquid nitrogen for 4 s, and the whole brain was removed and sectioned coronally (∼1 mm) on ice using a mouse brain slicer matrix (Brain Science Idea Co., Ltd., Osaka, Japan). The NAc was collected using a biopsy punch (Kai corporation, Tokyo, Japan; diameter 1.5 mm) on an ice-cold plate. The tissue samples were snap frozen in liquid nitrogen and stored at −80°C until use.

#### Immunoblotting

Sodium dodecyl sulphate-polyacrylamide gel electrophoresis (SDS-PAGE) was performed using a 6%, 8%, or 10% polyacrylamide gel (Nacalai Tesque). Proteins were separated using SDS-PAGE and transferred to polyvinylidene difluoride membranes (Immobilon-FL, Merck). The membranes were blocked for 30 min with Blocking-One or Blocking-One P (Nacalai Tesque) and incubated with primary antibodies for 1 h at room temperature (RT; 23–25°C) or overnight at 4°C. After washing, the membranes were incubated for 30 min with goat anti-rabbit IgG Alexa Fluor 680 (RRID: AB_2535758, Thermo Fisher Scientific), goat anti-mouse IgG Alexa Fluor 680 (RRID: AB_2535724, Thermo Fisher Scientific), or anti-mouse IgG DyLight 800 Conjugate (RRID: AB_10693543, Cell Signaling Technology) at RT for 30 min. Specific binding was detected using an infrared imaging system (LI-COR Biosciences, Lincoln, NE, USA). Band intensities were quantified using the ImageStudio software (RRID: SCR_015795, LI-COR Biosciences).

#### In vitro phosphorylation assay

The phosphorylation assay was performed as described previously^64^. ARHGEF2, ARHGAP21, and SHANK3 fragments were expressed in *E. coli* cells as GST fusion proteins and purified using glutathione-sepharose 4B beads. The kinase reaction for CaMKII was performed in 100 µL of a reaction mixture (25 mM Tris/HCl [pH 7.5], 2 mM Ca^2+^, 0.2 µM calmodulin, 1 mM DTT, 5 mM MgCl_2_, 50 µM [γ-^32^ P] ATP [1–20 GBq/mmol], 0.05 µM CaMKII, and 0.3 µM purified GST-proteins) for 30 min at 30°C. The kinase reaction for Rho-kinase was performed in 100 µL of a reaction mixture (25 mM Tris/HCl [pH 7.5], 1 mM EDTA, 1 mM ethylene glycol tetraacetic acid (EGTA), 1 mM DTT, 5 mM MgCl_2_, 50 µM [γ-^32^ P] ATP [1–20 GBq/mmol], 0.05 µM Rho-kinase-cat, and 0.3 µM purified GST-proteins) for 30 min at 30°C. The reaction mixtures were boiled in SDS sample buffer and subjected to SDS-PAGE and silver staining. Radiolabelled proteins were visualised using an image analyser (FLA9000; GE Healthcare).

#### Mass spectrometry

Mass spectrometry was performed as described previously with some modifications^13^. Frozen mouse brain slices were lysed in lysis buffer (50 mM Tris/HCl, 1 mM EDTA, 150 mM NaCl, 1% NP-40, 0.5% sodium deoxycholate, 0.1% SDS, protease inhibitor cocktail [Roche], and PhosStop [Roche]; pH 7.5) and sonicated for 30 s. The lysate was centrifuged at 20,000 ×*g* at 4°C for 10 min. Subsequently, the lysates were adjusted to equal protein concentrations and incubated with 500 pmol of GST-14-3-3 immobilised on glutathione-sepharose 4B beads (GE Healthcare) for 60 min at 4°C. The beads were washed three times with lysis buffer and three times with wash buffer (50 mM Tris/HCl, 1 mM EDTA, and 150 mM NaCl; pH 7.5).

The bound proteins were extracted from the beads using guanidine solution (7 M guanidine and 50 mM Tris), reduced via incubation in 5 mM dithiothreitol for 30 min, and alkylated using 10 mM iodoacetamide for 1 h in the dark. The proteins were demineralised, concentrated via methanol/chloroform precipitation, and digested with trypsin (50 mM NH_4_HCO_3_, 1.2 M urea, and 0.5 μg trypsin). Phosphopeptide enrichment was performed using a Titansphere Phos-TiO Kit (GL Sciences, Tokyo, Japan), followed by demineralisation using SPE c-tips (Nikkyo Technos, Tokyo, Japan) according to the manufacturer’s instructions.

The peptides were analysed using liquid chromatography/mass spectrometry (MS) using with an Orbitrap Fusion mass spectrometer (Thermo Fisher Scientific) coupled to an UltiMate3000 RSLCnano LC system (Dionex Co., Amsterdam, Netherlands) using a nano HPLC capillary column (Nikkyo Technos Co.; internal diameter, 75 μm; length, 150 mm; particle size, 3 μm) via a nanoelectrospray ion source.

Reversed-phase chromatography was performed using a linear gradient (0 min, 5% B; 100 min, 40% B) of solvent A (2% acetonitrile with 0.1% formic acid) and solvent B (95% acetonitrile with 0.1% formic acid) at an estimated flow rate of 300 nL/min. Prior to MS/MS analysis, a precursor ion scan was carried out using a 400–1600 mass-to-charge ratio (m/z). Tandem MS was performed by isolation at 0.8 Th with a quadrupole, HCD fragmentation with a normalised collision energy of 30%, and rapid-scanning MS analysis in the ion trap. Only the precursors with charge states of 2–6 were sampled for MS2. The dynamic exclusion duration was set to 10 s with 10 ppm tolerance. The instrument was operated in the top-speed mode with 3 s cycles.

A peak list was generated and calibrated using the MaxQuant software (version 1.4.1.2). The reference proteome of *Mus musculus* was obtained from UniProtKB in March 2023, and database searches against this genome were performed using the MaxQuant software. The carbamidomethylation of cysteine was set as a fixed modification, and the oxidation of methionine, phosphorylation of Ser/Thr/Tyr, and N-terminal acetylation were set as variable modifications. The false discovery rates for the peptide, protein, and site levels were set to 0.01. The ion peak intensities obtained from over three independent experiments were analysed. Phosphoproteins and their phosphorylation sites were stimulated more than two-fold than those in the control and were detected at least twice in over three independent experiments.

The identified proteins were considered candidate substrates for Rho-kinase if they fulfilled the following three criteria: the phosphorylation level of the protein based on the ion intensity was increased more than two-fold by treatment with CLA and detected at least twice in over three independent experiments; the phosphorylation of the protein was decreased by treatment with Y-27632; and the phosphorylation motif was consistent with the consensus for Rho-kinase [(R/K)XX(pS/pT)] or [(R/K)X(pS/pT)].

#### GST-RhoA (G17A) pulldown assay

The guanine nucleotide exchange activity of ARHGEF2 was measured as described previously, with some modifications^28^. Wild-type, T103E mutant, or T103A mutant ARHGEF2; and GFP-tagged wild-type, constitutively active (CA), or dominant negative (DN) CaMKII were transfected into COS-7 cells using the Lipofectamine 2000 transfection reagent (Thermo Fisher Scientific). COS-7 cells were lysed in 20 mM HEPES (pH 7.5), 1 mM dithiothreitol, 150 mM NaCl, 1 mM EDTA, 1% Triton X-100, protease inhibitor cocktail (Roche), and PhosStop (Roche). Cell lysates were obtained via centrifugation at 20,000 ×*g* for 10 min at 4°C and were incubated with Purified GST-RhoA (G17A)-glutathione magnetic beads (Thermo Fisher Scientific) for 1 h at 4°C. Protein complexes were dissolved in the SDS-PAGE sample buffer, and the eluted samples were analysed via immunoblotting.

#### GST-Rhotekin-RBD pulldown assay

RhoA activity was determined using a pull-down assay with the GST-Rho–binding domain of Rhotekin (GST-Rhotekin-RBD) as described previously^65^. Briefly, the striatal/accumbal slices were sonicated in 300 µL lysis buffer (50 mM Tris-HCl [pH 7.5], 1 mM EGTA, 20 mM MgCl_2_, 500 mM NaCl, 0.5% NP-40, protease inhibitor cocktail [Roche], and PhosStop [Roche] containing 20 μg of GST-Rhotekin-RBD). The lysates were centrifuged at 20,000 ×*g* for 3 min at 4°C, and the supernatants were incubated with glutathione-coated magnetic beads (Thermo Fisher Scientific) for 30 min at 4°C. The beads were washed with an excess of lysis buffer and then eluted with SDS-sample buffer. The eluates were subjected to SDS-PAGE, followed by immunoblot analysis with the anti-RhoA antibody.

#### Immunofluorescence analysis

Mice were anesthetised through intraperitoneal administration of a mixture of 0.75 mg/kg of medetomidine (Domitor, Nippon Zenyaku Kogyo, Japan), 4.0 mg/kg of midazolam (Dormicum, Maruishi Pharmaceutical, Osaka, Japan), and 5.0 mg/kg of butorphanol (Vetorphale, Meiji Seika Pharma, Osaka, Japan). They were then transcardially perfused with 4% paraformaldehyde. The brains were removed, incubated in 4% paraformaldehyde overnight at 4°C, and sectioned coronally (thickness, 50–100 μm) using a vibratome (VT1200S, Leica Microsystems). Brain slices were incubated with 0.1% Triton X-100/phosphate-buffered saline for 10 min and blocked with Blocking-One P for 30 min. Brain slices and neurons were incubated with the indicated antibodies overnight at 4°C. After washing, the samples were incubated with the Alexa

Fluor 405-, 488-, or 555-conjugated secondary antibody for 1 h. The nuclei were visualised by staining with TO-PRO-3 iodide (Thermo Fisher Scientific). Confocal images were recorded using LSM 780 microscopes built around an Axio Observer Z1 with Plan-Apochromat 20× (numerical aperture [NA] 0.8), a C-Apochromat 40× (NA 1.2), or a Plan-Apochromat 63× (NA 1.40) lens controlled using the ZEN Digital Imaging for Light Microscopy software (RRID: SCR_013672, Carl Zeiss, Oberkochen. Germany). Confocal images for dendritic spine were obtained using LSM 980 microscopes built around an Axio Observer 7 with Plan Apochromat 63xOil Airy 63× (NA 1.40) lens controlled by the ZEN Digital Imaging for Light Microscopy software (Carl Zeiss). The entire coronal brain section was observed using fluorescence microscopes (BZ-X800, Keyence, Osaka, Japan).

#### Adeno-associated virus preparation

Adeno-associated virus (AAV) vectors were prepared and titered as described previously^66^. Briefly, plasmids for the AAV vector, pHelper (Cell BioLabs) and pAAV-DJ (Cell BioLabs) were transfected into AAV293 cells. After 3 days of incubation, the cells were harvested, and the AAV vectors produced in AAV293 cells were purified by centrifugation using CsCl-gradient ultracentrifugation. CsCl was removed via dialysis, and the AAV titres were estimated using quantitative polymerase chain reaction.

#### AAV injection

Mice were anesthetised with a mixture of medetomidine, midazolam, and butorphanol and then positioned in a stereotaxic frame (David Kopf, Tujunga, CA, USA) preoperatively. Postoperatively, mice were intraperitoneally administered 0.75 mg/kg atipamezole (Antisedan; Nippon Zenyaku Kogyo) to reverse the anesthesia. An AAV vector (0.5 μL, 1.0×10^12^ genome copies/mL) was injected into the bilateral NAc through a glass microinjection capillary tube at a 10° angle at a rate of 0.1 μL/min (0.5 μL/site; four sites). AAV vectors were infused unilaterally in the NAc at two dorsal-ventral levels. The anteroposterior (AP), mediolateral (ML), and dorsoventral (DV) coordinates relative to the bregma were as follows (in mm): (1) AP +1.6, ML +1.5, and DV −4.4, respectively, and (2) AP +1.0, ML +1.6, and DV −4.5, respectively. For Ca^2+^-imaging using wireless fiber photometry, AAV-Flex-GCaMP6 were infused in the NAc (AP +1.3, ML +1.5, and DV −4.5) of Arora2a-Cre mice. The TeleFipho cannula (core 400 μm, NA 0.39, cladding 425 mm, ferrule 2.5 mm) was implanted at the same stereotaxic coordinates. Cannula was fixed to the skull with light-cured bonding adhesive (Kuraray Noritake Dental Inc, Japan), and the exposed skull was covered with light-cured composite resin (Kuraray Noritake Dental Inc, Japan). Animals were allowed to recover for 2 weeks before the beginning of handling and Ca^2+^-imaging experiments.

#### Behavioural analysis

A step-through passive avoidance test was conducted as described previously^51^. The experimental apparatus consisted of two compartments (one illuminated and one dark; volume: 25 × 15 × 15 cm^3^; height, 15 cm) equipped with a grid floor. The two compartments were separated with a guillotine door. The entire test consisted of three sessions conducted over 3 consecutive days (24 h apart) as follows. (1) During the habituation phase, the mouse was placed in the illuminated compartment. After 15 s, the door was opened, and the mouse was given 2 min to enter the dark compartment. Once the mouse entered the dark compartment, the door was closed, and the animal was removed. (2) During the training phase, the mouse was placed in the illuminated compartment with the door closed. After 15 s, the door was opened, and the mouse was allowed to enter the dark compartment. Once the mouse entered the dark compartment (with all four paws in), the door was closed, and an electric foot shock (0.7 mA; duration, 3 s) was delivered through the grid floor. The mouse was returned to its cage after 15 s. (3) In the testing phase, the mouse was again placed in the illuminated compartment, and the door was opened after 15 s. The latency to enter the dark compartment was recorded for up to 300 s. When the mice did not enter the room for at least 300 s, a score of 300 s was assigned.

#### Calcium imaging in freely moving mice

*In vivo* Calcium imaging was performed using a TeleFipho wireless fiber photometry device (Bioresearch Center Inc., Aichi, Japan). Beginning 2 weeks after injection of the AAV, a dummy head stage was attached to the implanted cannula, and mice were kept in their cages for 3 days. Saline or MK-801 (0.1 mg/kg, s.c.) was administered to the mouse 30 min before calcium imaging. After the dummy head stage was attached to the mouse and the baseline of calcium activity had stabilized, the mouse was placed in the illuminated compartment with the door closed. After 15 s, the door was opened, and the mouse was allowed to enter the dark compartment. Once the mouse entered the dark compartment, the door was closed, and an electric foot shock (0.7 mA; duration, 3 s) was delivered through the grid floor. Calcium activity during electric foot shock was continuously recorded. Data were normalised against the mean of 2 s baseline (F) before the electric foot shock (time=0). Changes in fluorescence following the electric shock were quantified by measuring the area under the ΔF/F curve. Heatmaps were generated using the R software (version 4.0.2).

#### Dendritic spine imaging and analysis

Dendritic spine imaging was performed as previously described^51^. Three weeks after mice were injected with AAV virus, mice were transcardially perfused with 4% PFA in PBS. The whole brains were removed and immersed in 4% PFA in PBS overnight at 4 C to postfix the brains. The brains were sectioned coronally (100 μm) by using a VT1200S vibratome (Leica Microsystems, Wetzlar, Germany). The slices were then immunostained with anti-GFP antibody. For spine morphology analysis, Z-stacks of secondary or tertiary dendrites (>20 μm long, >50 μm away from the soma) were captured with an LSM 980 confocal microscope using a ×63 objective (NA = 1.40, 5X zoom). All dendrites and spines within images were traced using Imaris Filament Tracer (RRID: SCR_007366, Bitplane, Switzerland) and analyzed using Imaris software (version 9.8.2, RRID: SCR_007370). Spines were subsequently classified with the following morphological criteria as described previously^67^ with a minor modification: mushroom spines were identified as spines with maximal width (head) > 0.6μm and maximal width (head)/mean width (neck) >1.1; stubby spines were identified as spines with maximal width (head)/mean width (neck) <1.1 and length (spine)/ maximal width (head) <2.5; thin spines were the remaining spines.

#### In vitro electrophysiology

Mice were transcardially perfused under isoflurane anesthesia with an ice-cold dissection buffer containing (in mM) 87 NaCl, 25 NaHCO_3_, 25 D-glucose, 2.5 KCl, 1.25 NaH_2_PO_4_, 0.5 CaCl_2_, 7 MgCl_2_, and 75 sucrose, aerated with 95% O_2_C+C5% CO_2_. The mice were then decapitated, and the brain was isolated and cut into coronal slices (250–300 μm thick) on a vibratome in the ice-cold dissection buffer. The slices containing the NAc were incubated for 30 min at 35°C in the dissection buffer and maintained thereafter at RT in an artificial cerebrospinal fluid (aCSF) containing (in mM) 125 NaCl, 25 NaHCO_3_, 25 D-glucose, 2.5 KCl, 1.25 NaH_2_PO_4_, 1 MgCl_2_, and 2 CaCl_2_), aerated with 95% O_2_ and 5% CO_2_. The ROCK inhibitor Y-27632 was stocked in DMSO and diluted into the aCSF so that the final concentration of DMSO was 0.21%. For control experiments, 0.21% DMSO was added to the aCSF.

Whole-cell patch-clamp recordings were performed using an IPA amplifier (Sutter Instruments) at RT. Fluorescently labelled D2R-MSNs were visually identified using an upright microscope (BX51WI; Olympus) equipped with a scientific complementary metal-oxide-semiconductor video camera (Zyla4.2plus; Andor). The recording pipettes were filled with the intracellular solution containing (in mM): 135 potassium gluconate, 4 KCl, 4 Mg-ATP, 10 Na_2_-phosphocreatine, 0.3 Na-GTP, and 10 HEPES (pH 7.3, 295CmOsm). Patch pipettes (5–7CMΩ) had a series resistance of 6.5–30CMΩ. Data were filtered at 5 kHz, digitised at 10 kHz, and recorded using the SutterPatch software running on Igor Pro 8. Cells with a depolarised resting membrane potential greater than –70 mV were not used for experiments. Excitatory postsynaptic potentials (EPSPs) were evoked at 0.05 Hz by focal extracellular stimulation with an aCSF-filled glass electrode positioned at ∼150–200 μm from the recording pipette. The intensity for the extracellular stimulation (0.2 ms, 5–30 μA) was adjusted to evoke single-component EPSPs. After 10 minutes of baseline recording, pairing stimuli were applied to induce spike-timing-dependent synaptic plasticity (STDP). Backpropagating action potentials (bAPs) were induced by direct somatic current injection (5 ms, 1.5 nA). The pairing protocol (STDP protocol) was modified from that of Shen et al. (2008)^68^ and consisted of 30 trains of five bAP bursts repeated at 0.1 Hz, with each bAP burst composed of four bAPs preceded by four EPSPs (+5 ms) evoked at 50 Hz. During recordings, a GABAA receptor antagonist (gabazine, 10 μM) was routinely added to the aCSF to record EPSPs, and an A2A receptor agonist (CGS21680, 0.3 μM) was also routinely bath applied to stably induce long-term potentiation in the control condition.

### QUANTIFICATION AND STATISTICAL ANALYSES

Data analysis was performed using the Prism 9 statistics software (RRID: SCR_002798; GraphPad Software, Inc., La Jolla, CA, USA). All data are expressed as the mean ±standard error of the mean (SEM). A one-way or two-way analysis of variance was performed, followed by Tukey’s or Dunnett’s multiple-comparison test when the F ratios were significant (p<0.05). The paired t-test, Unpaired t test, and Mann–Whitney U test determined significant differences between the two groups. In the passive avoidance tests, the Kruskal–Wallis H test followed by Dunn’s multiple-comparison test or the Mann–Whitney U test was used because the distribution was skewed due to the existence of a cut-off time. The calculated values are shown in Supplementary Table 3.

### Data and code availability

#### Lead contact

Further information and requests for resources and reagents should be directed to and will be fulfilled by the lead contact, Kozo Kaibuchi.

#### Materials availability

All unique/stable reagents generated in this study will be made available on request; however, we may require a payment and/or a completed Materials Transfer Agreement if there is a potential for commercial application.

#### Data and code availability

All data are available in Dryad data repository (https://doi.org/10.5061/dryad.fttdz08zr) Relevant datasets for Table 1, Supplementary Table 1 and Supplementary Table 2 in the paper are available in KANPHOS database (https://kanphos.neuroinf.jp).

## Acknowledgments

We are grateful to Dr. K. Kobayashi for providing us the Drd2-YFP transgenic mice and to Dr. Young-Bum Kim (Harvard Medical School, USA) for providing us the Rock1 floxed mice. We are grateful Dr. C. Yokoyama for invaluable comments on our manuscript. We are grateful to M, Fukushima, M. Taguchi, A. Hara, S. Tanaka and other Kaibuchi lab members for helpful discussions and preparation of some materials, K. Taki for help with MS data acquisition, and T. Ishii for secretarial assistance. We also thank the Division for Research on Laboratory Animals, Radioisotope Center Medical Branch, Medical Research Engineering of Nagoya University Graduate School of Medicine, and Education and Research Center of Animal Models for Human Diseases at the Fujita Health University. This work was supported by the following funding sources: AMED, Grant Numbers JP19dm0207075 (K.K.) and JP21wm0425017 (T.N. and Y.F.); JSPS KAKENHI, Grant Numbers JP17H01380 (K.K.), JP18K14849 (Y.F.), JP21K06428 (Y.F.), and JP21K06427 (D.T.); Uehara Science Foundation (K.K. and Y.F.), Takeda Science Foundation (K.K. and Y.F.), and Hori Sciences and Arts Foundation (K.K. and Y.F.). This work was also supported by the Collaborative Research Project (2011-2302) of the Brain Research Institute of Niigata University.

## Author contributions

Conceptualisation, Y.F. and K.K.; Methodology, D.T., T.N., M.A. and T.Y; Formal analysis and investigation, Y.F., R.U.A., X.Z., E.H., M.K., S.N., M.W., X.L., H.W., O.F., H.S. Y.L., M.A., A.Y., and Y.N.; Writing - original draft preparation, Y.F. and R.U.A.; Writing - review and editing, Y.F. and K.K.; Funding acquisition, Y.F., D.T., T.N., and K.K.; Resources, K.Y., S.U., and K.S.; Supervision, K.K.; Project administration, K.K. All authors approved the final version of the manuscript.

## Additional Information

- Supplementary Information is available for this paper.
- Correspondence and requests for materials should be addressed to Kozo Kaibuchi.
- Peer review information:
- Reprints and permissions information is available at www.nature.com/reprints.

## Supplementary information

### Supplementary Figures

**Supplementary Fig. 1.**
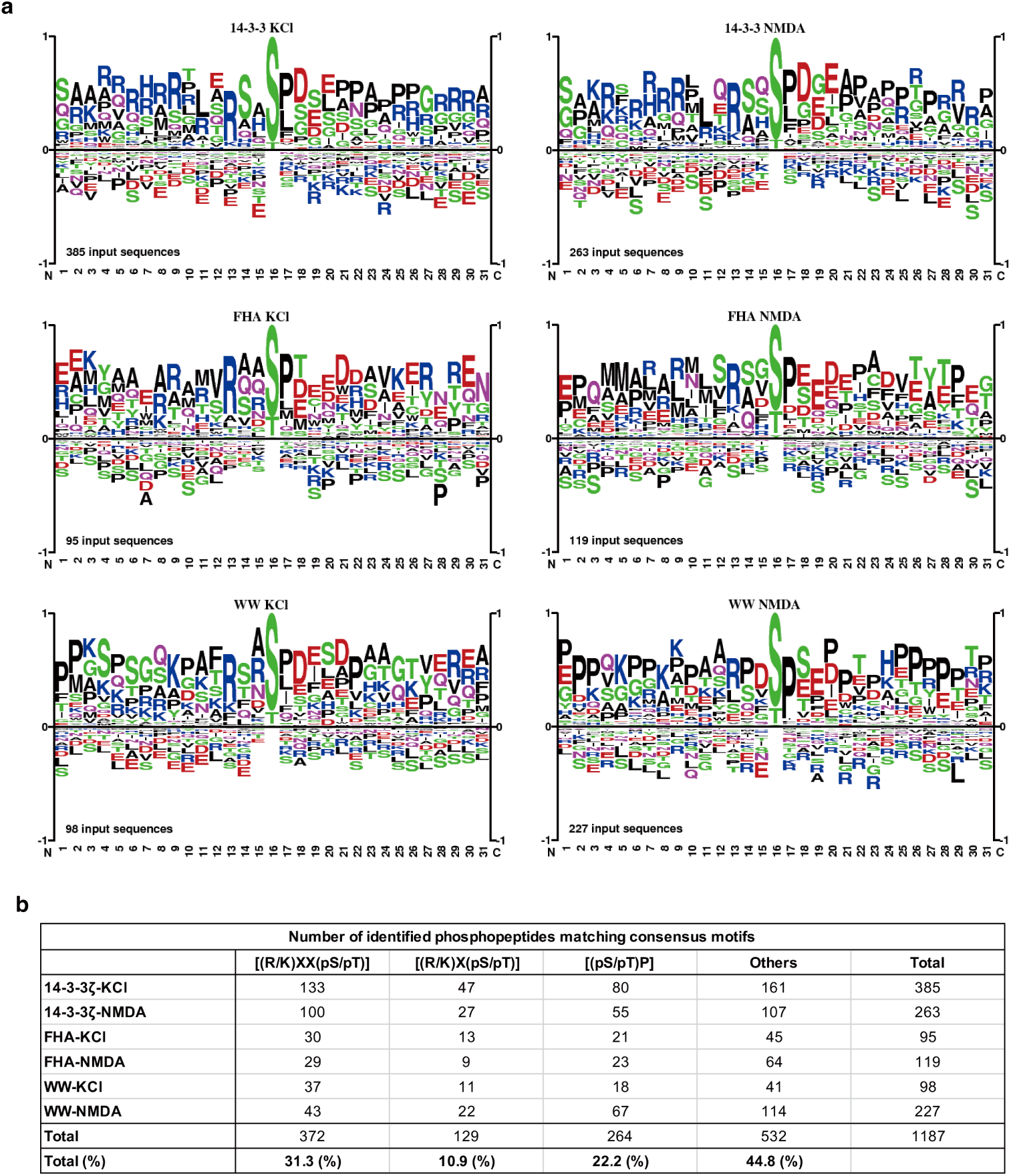
related to Figure 1. Sequence motif analysis of identified peptides by phosphoproteomic analysis of High K^+^ and NMDA. **(a)** The sequence alignment of the phosphopeptides bound to each of the WW, FHA, and 14-3-3ζ baits, showing different phosphopeptide binding preferences (http://www.phosphosite.org) is shown. **(b)** Summary of sequencing motif analysis using identified peptides.

**Supplementary Fig. 2.**
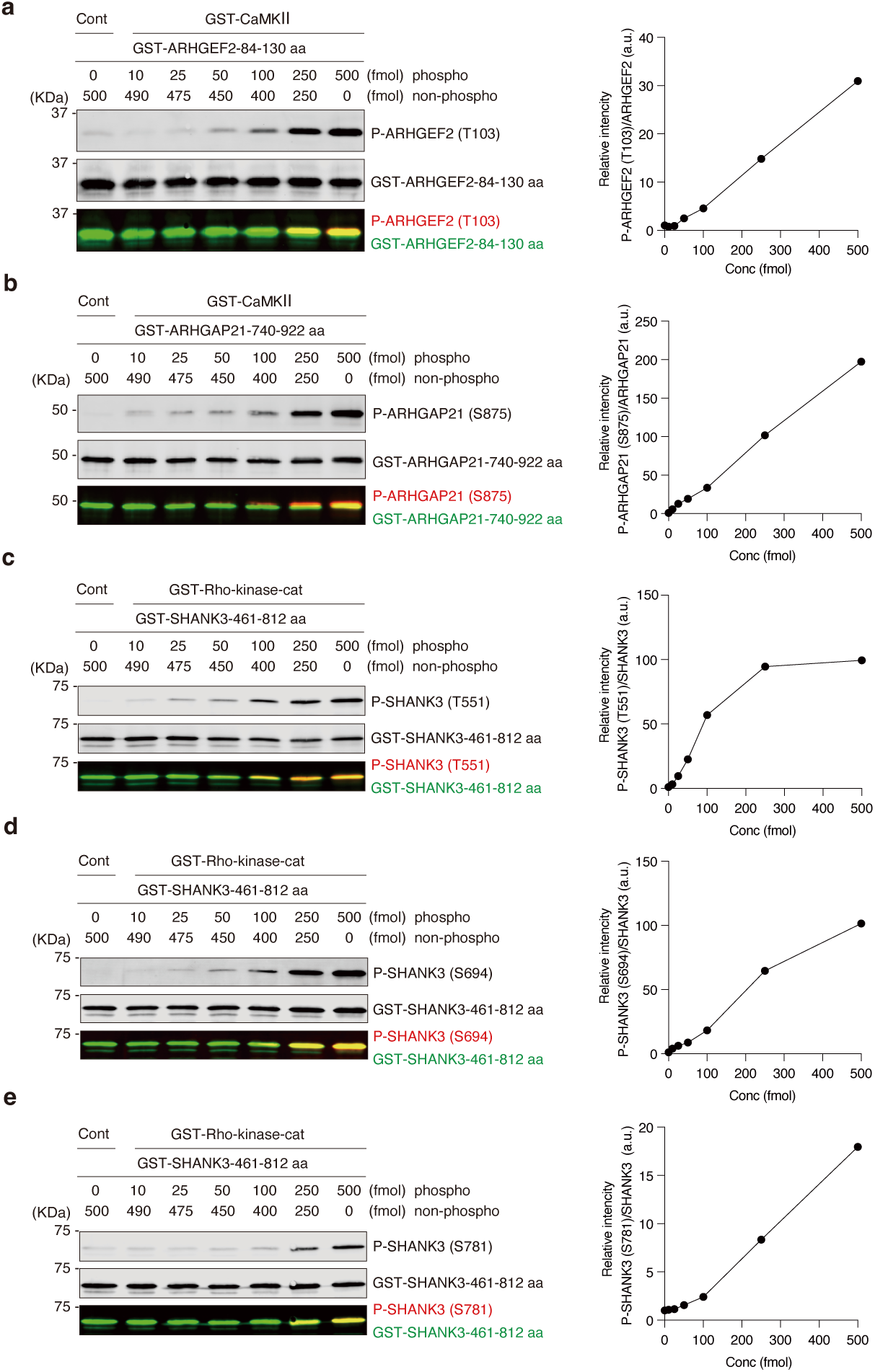
related to Figure 2, 3 and 5. Specificity of generated phospho-specific antibodies. The specificity of the antibodies against ARHGEF2 at T103 **(a)**; ARHGAP21 at S875 **(b)**; and SHANK3 at T551 **(c)**, S694 **(d)**, or S781 **(e)**. For SDS-PAGE and immunoblot analysis with the indicated antibodies, 500 fmol of GST-proteins containing the indicated amount of phosphorylated and unphosphorylated proteins is used.

**Supplementary Fig. 3.**
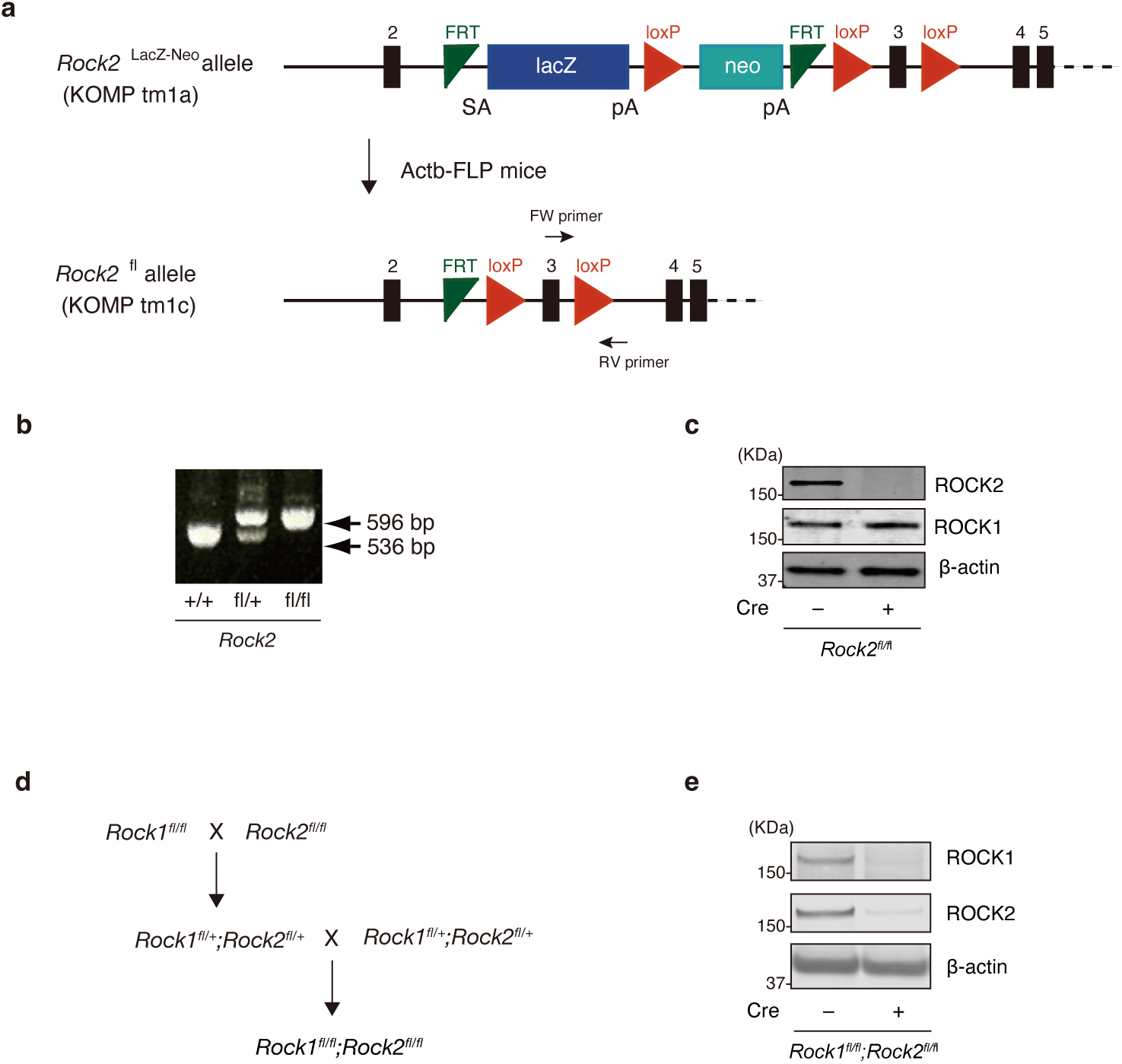
related to Figure 4. Generation of *Rock1/2* floxed mice. **(a)** Schematic representation of the initial allele (tm1a) containing LacZ reporter-promoter-driven neo targeting cassette, FLP-flippase recombinase target (FRT) sites, and loxP sites. The tm1a mice were crossed with Actb-Flp mice to generate *Rock2* floxed mice (tm1c). **(b)** Genotype verification of *Rock2* floxed mice by genomic PCR. **(c)** Cultured striatal neurons obtained from homozygous *Rock2* floxed mice (*Rock2*^fl/fl^) were infected with AAV-CAGGS-Cre at DIV4. At 14 days after infection, immunoblot analysis was performed with anti-ROCK1, anti-ROCK2, and beta-actin antibodies. **(d)** Schematic representation of generation of *Rock1* and *Rock2* double-floxed mice. Heterozygous *Rock1* and *Rock2* double floxed mice (*Rock1*^fl/+^*;Rock2*^fl/+^) were obtained by crossing homozygous *Rock1* floxed mice (*Rock1*^fl/fl^) with homozygous *Rock2* floxed mice (*Rock2*^fl/fl^). **(e)** Cultured striatal neurons obtained from homozygous *Rock1* and *Rock2* double floxed mice (*Rock1*^fl/fl^*;Rock2*^fl/fl^) were infected with AAV-CAGGS-Cre at DIV7. At 14 days after infection, immunoblot analysis was performed with anti-ROCK1, anti-ROCK2, and beta-actin antibodies.

**Supplementary Fig. 4.**
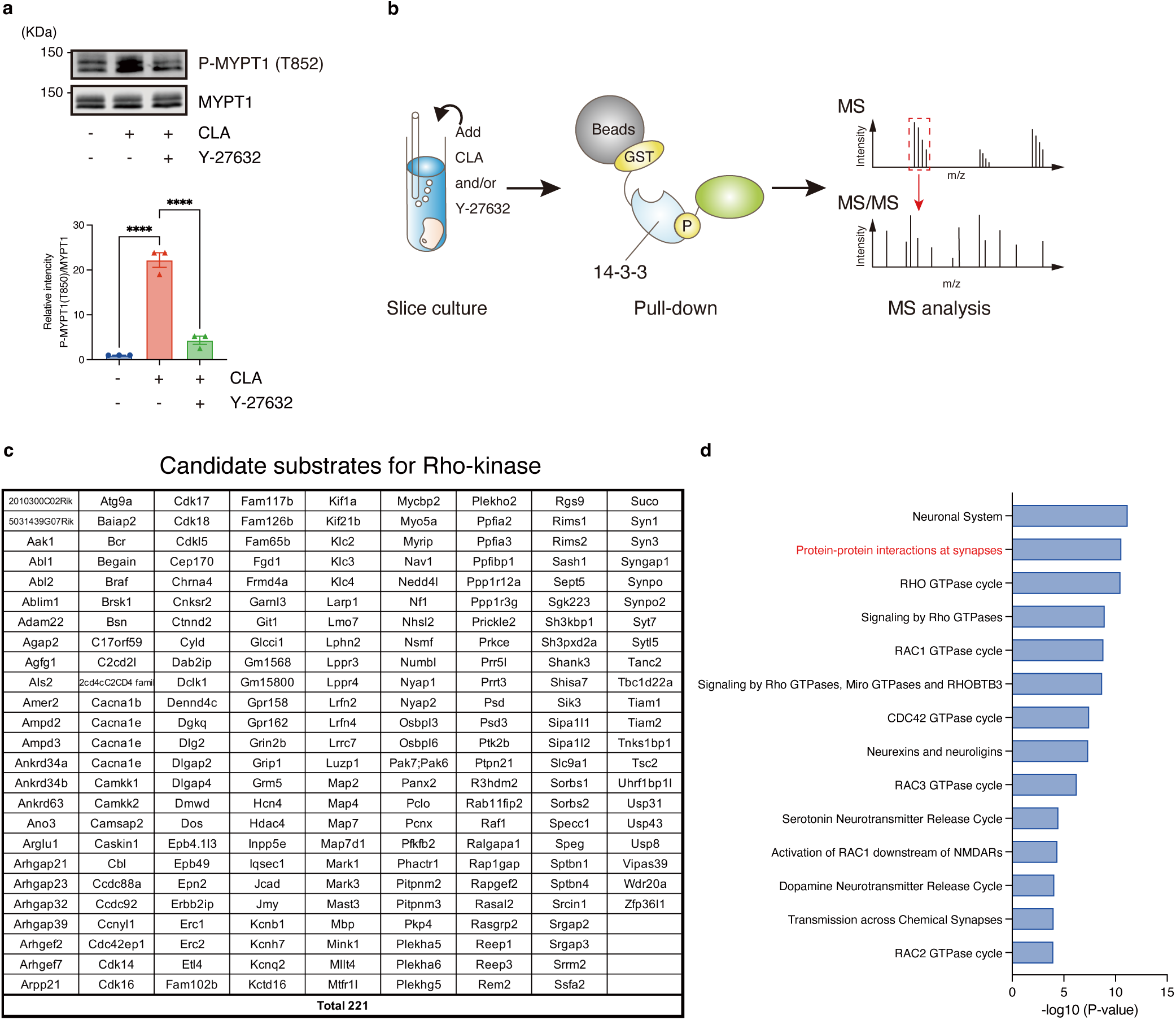
related to Figure 5. Screening of candidate substrates for Rho-kinase. **(a)** Treatment with calyculin-A (350 nM) for 60 min induced phosphorylation of MYPT1 at T852, whereas treatment with calyculin-A (350 nM) for 60 min and Y-27632 (20 µM) for 60 min reduced the level of phosphorylation. Data present the mean ± SEM of three independent experiments. One-way ANOVA followed by Tukey’s multiple comparisons test. ****p<0.0001. **(b)** Schematic representation of the Kinase-Oriented Substrate Screening (KIOSS) method. Extracts of these striatal/accumbal slices were applied to affinity beads coated with 14-3-3 ζ to enrich the phosphoproteins. The bound proteins were digested using trypsin and subjected to LC-MS/MS to identify the phosphorylated proteins and their phosphorylation sites. **(c)** List of candidate substrates for Rho-kinase. **(d)** Pathway analysis using the Reactome database (available online at: http://www.reactome.org) of 221 proteins.

**Supplementary Fig. 5.**
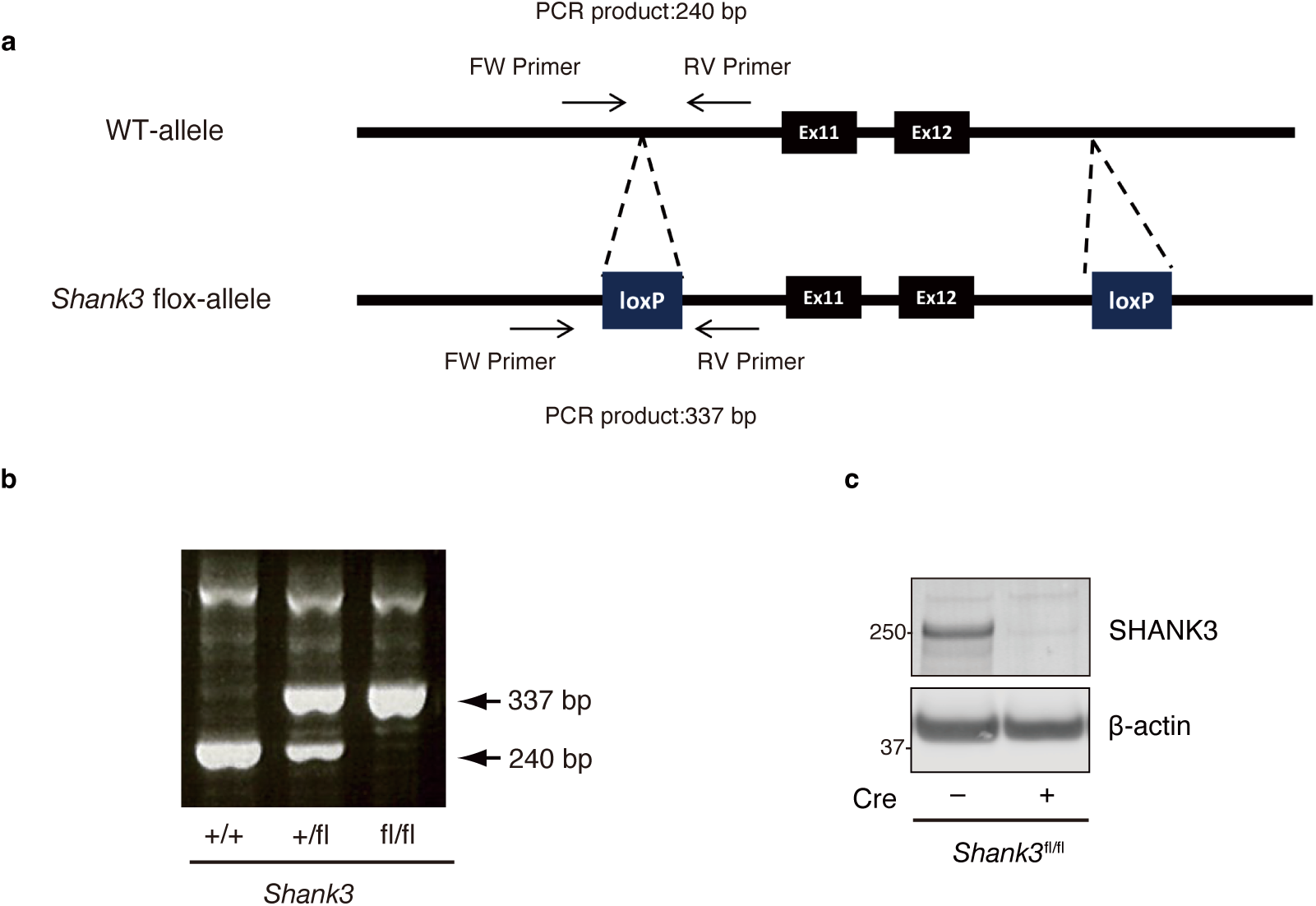
related to Figure 6. Generation of *Shank3* floxed mice. **(a)** Schematic representation of *Shank3* floxed mice. A DNA fragment containing a loxP sequence and Pgk1 promoter-driven Neo flanked by a pair of FRT sequences was inserted upstream of exon 12. The other loxP site was introduced downstream of exon 11. **(b)** Genotype verification of *Shank3* floxed mice by genomic PCR. **(c)** Cultured striatal neurons obtained from homozygous *Shank3* floxed mice (*Shank3*^fl/fl^) were infected with AAV-CAGGS-Cre at DIV4. At 14 days after infection, immunoblot analysis was performed with anti-SHANK3 and beta-actin antibodies.

### Supplementary Tables

**Supplementary Table 1. Detailed information of phosphoproteomic analysis of KCl and NMDA.**

**Supplementary Table 2. Detailed information of phosphoproteomic analysis of Rho-kinase.**

**Supplementary Table 3. Summary of statistical analysis in this study.**

## References

1. Lamprecht R, LeDoux J. Structural plasticity and memory. Nat Rev Neurosci 5, 45–54 (2004).

2. Luscher C, Malenka RC. NMDA receptor-dependent long-term potentiation and long-term depression (LTP/LTD). Cold Spring Harb Perspect Biol 4, (2012).

3. Paoletti P, Bellone C, Zhou Q. NMDA receptor subunit diversity: impact on receptor properties, synaptic plasticity and disease. Nat Rev Neurosci 14, 383–400 (2013).

4. Lisman J, Schulman H, Cline H. The molecular basis of CaMKII function in synaptic and behavioural memory. Nat Rev Neurosci 3, 175–190 (2002).

5. Nicoll RA, Schulman H. Synaptic memory and CaMKII. Physiol Rev 103, 2877–2925 (2023).

6. Barria A, Derkach V, Soderling T. Identification of the Ca2+/calmodulin-dependent protein kinase II regulatory phosphorylation site in the alpha-amino-3-hydroxyl-5-methyl-4-isoxazole-propionate-type glutamate receptor. J Biol Chem 272, 32727–32730 (1997).

7. Strack S, McNeill RB, Colbran RJ. Mechanism and regulation of calcium/calmodulin-dependent protein kinase II targeting to the NR2B subunit of the N-methyl-D-aspartate receptor. J Biol Chem 275, 23798–23806 (2000).

8. Mammen AL, Kameyama K, Roche KW, Huganir RL. Phosphorylation of the alpha-amino-3-hydroxy-5-methylisoxazole4-propionic acid receptor GluR1 subunit by calcium/calmodulin-dependent kinase II. J Biol Chem 272, 32528–32533 (1997).

9. Lammel S, et al. Input-specific control of reward and aversion in the ventral tegmental area. Nature 491, 212–217 (2012).

10. Hikida T, Kimura K, Wada N, Funabiki K, Nakanishi S. Distinct roles of synaptic transmission in direct and indirect striatal pathways to reward and aversive behavior. Neuron 66, 896–907 (2010).

11. Lim SA, Kang UJ, McGehee DS. Striatal cholinergic interneuron regulation and circuit effects. Front Synaptic Neurosci 6, 22 (2014).

12. Nakanishi S, Hikida T, Yawata S. Distinct dopaminergic control of the direct and indirect pathways in reward-based and avoidance learning behaviors. Neuroscience 282, 49–59 (2014).

13. Nagai T, et al. Phosphoproteomics of the Dopamine Pathway Enables Discovery of Rap1 Activation as a Reward Signal In Vivo. Neuron 89, 550–565 (2016).

14. Nishioka T, Shohag MH, Amano M, Kaibuchi K. Developing novel methods to search for substrates of protein kinases such as Rho-kinase. Biochim Biophys Acta 1854, 1663–1666 (2015).

15. Shohag MH, et al. Phosphoproteomic Analysis Using the WW and FHA Domains as Biological Filters. Cell Struct Funct 40, 95–104 (2015).

16. Tsuboi D, et al. Dopamine drives neuronal excitability via KCNQ channel phosphorylation for reward behavior. Cell Rep 40, 111309 (2022).

17. Nishioka T, Nakayama M, Amano M, Kaibuchi K. Proteomic screening for Rho-kinase substrates by combining kinase and phosphatase inhibitors with 14-3-3zeta affinity chromatography. Cell Struct Funct 37, 39–48 (2012).

18. Ahammad RU, et al. KANPHOS: A Database of Kinase-Associated Neural Protein Phosphorylation in the Brain. Cells 11, (2021).

19. Rust HL, Thompson PR. Kinase consensus sequences: a breeding ground for crosstalk. ACS Chem Biol 6, 881–892 (2011).

20. Thomas GM, Huganir RL. MAPK cascade signalling and synaptic plasticity. Nat Rev Neurosci 5, 173–183 (2004).

21. Murakoshi H, Wang H, Yasuda R. Local, persistent activation of Rho GTPases during plasticity of single dendritic spines. Nature 472, 100–104 (2011).

22. Lazarini M, et al. ARHGAP21 is a RhoGAP for RhoA and RhoC with a role in proliferation and migration of prostate adenocarcinoma cells. Biochim Biophys Acta 1832, 365–374 (2013).

23. Ren Y, Li R, Zheng Y, Busch H. Cloning and characterization of GEF-H1, a microtubule-associated guanine nucleotide exchange factor for Rac and Rho GTPases. J Biol Chem 273, 34954–34960 (1998).

24. Bos JL, Rehmann H, Wittinghofer A. GEFs and GAPs: critical elements in the control of small G proteins. Cell 129, 865–877 (2007).

25. Peck J, Douglas Gt, Wu CH, Burbelo PD. Human RhoGAP domain-containing proteins: structure, function and evolutionary relationships. FEBS Lett 528, 27–34 (2002).

26. Govek EE, Newey SE, Van Aelst L. The role of the Rho GTPases in neuronal development. Genes Dev 19, 1–49 (2005).

27. Schmidt A, Hall A. Guanine nucleotide exchange factors for Rho GTPases: turning on the switch. Genes Dev 16, 1587–1609 (2002).

28. Garcia-Mata R, Wennerberg K, Arthur WT, Noren NK, Ellerbroek SM, Burridge K. Analysis of activated GAPs and GEFs in cell lysates. Methods Enzymol 406, 425–437 (2006).

29. Kimura K, et al. Regulation of myosin phosphatase by Rho and Rho-associated kinase (Rho-kinase). Science 273, 245–248 (1996).

30. Friedman DP, Aggleton JP, Saunders RC. Comparison of hippocampal, amygdala, and perirhinal projections to the nucleus accumbens: combined anterograde and retrograde tracing study in the Macaque brain. J Comp Neurol 450, 345–365 (2002).

31. Musazzi L, et al. Acute stress increases depolarization-evoked glutamate release in the rat prefrontal/frontal cortex: the dampening action of antidepressants. PLoS One 5, e8566 (2010).

32. Zhu Y, Wienecke CF, Nachtrab G, Chen X. A thalamic input to the nucleus accumbens mediates opiate dependence. Nature 530, 219–222 (2016).

33. Gong S, et al. Targeting Cre recombinase to specific neuron populations with bacterial artificial chromosome constructs. J Neurosci 27, 9817–9823 (2007).

34. Gong S, et al. A gene expression atlas of the central nervous system based on bacterial artificial chromosomes. Nature 425, 917–925 (2003).

35. Dash PK, Orsi SA, Moody M, Moore AN. A role for hippocampal Rho-ROCK pathway in long-term spatial memory. Biochem Biophys Res Commun 322, 893–898 (2004).

36. Lamprecht R, Farb CR, LeDoux JE. Fear memory formation involves p190 RhoGAP and ROCK proteins through a GRB2-mediated complex. Neuron 36, 727–738 (2002).

37. Mueller BK, Mack H, Teusch N. Rho kinase, a promising drug target for neurological disorders. Nat Rev Drug Discov 4, 387–398 (2005).

38. Gurkar AU, et al. Identification of ROCK1 kinase as a critical regulator of Beclin1-mediated autophagy during metabolic stress. Nat Commun 4, 2189 (2013).

39. Shen W, Flajolet M, Greengard P, Surmeier DJ. Dichotomous dopaminergic control of striatal synaptic plasticity. Science 321, 848–851 (2008).

40. Rex CS, et al. Different Rho GTPase-dependent signaling pathways initiate sequential steps in the consolidation of long-term potentiation. J Cell Biol 186, 85–97 (2009).

41. Krucker T, Siggins GR, Halpain S. Dynamic actin filaments are required for stable long-term potentiation (LTP) in area CA1 of the hippocampus. Proc Natl Acad Sci U S A 97, 6856–6861 (2000).

42. Coles CH, Bradke F. Coordinating neuronal actin-microtubule dynamics. Curr Biol 25, R677–691 (2015).

43. Xu X, et al. 14-3-3zeta deficient mice in the BALB/c background display behavioural and anatomical defects associated with neurodevelopmental disorders. Sci Rep 5, 12434 (2015).

44. Sheng M, Hoogenraad CC. The postsynaptic architecture of excitatory synapses: a more quantitative view. Annu Rev Biochem 76, 823–847 (2007).

45. Naisbitt S, et al. Shank, a novel family of postsynaptic density proteins that binds to the NMDA receptor/PSD-95/GKAP complex and cortactin. Neuron 23, 569–582 (1999).

46. MacGillavry HD, Kerr JM, Kassner J, Frost NA, Blanpied TA. Shank-cortactin interactions control actin dynamics to maintain flexibility of neuronal spines and synapses. Eur J Neurosci 43, 179–193 (2016).

47. Wang X, et al. Synaptic dysfunction and abnormal behaviors in mice lacking major isoforms of Shank3. Hum Mol Genet 20, 3093–3108 (2011).

48. Yang M, et al. Reduced excitatory neurotransmission and mild autism-relevant phenotypes in adolescent Shank3 null mutant mice. J Neurosci 32, 6525–6541 (2012).

49. Durand CM, et al. SHANK3 mutations identified in autism lead to modification of dendritic spine morphology via an actin-dependent mechanism. Mol Psychiatry 17, 71–84 (2012).

50. Saneyoshi T, et al. Reciprocal Activation within a Kinase-Effector Complex Underlying Persistence of Structural LTP. Neuron 102, 1199–1210 e1196 (2019).

51. Yamahashi Y, et al. Phosphoproteomic of the acetylcholine pathway enables discovery of the PKC-beta-PIX-Rac1-PAK cascade as a stimulatory signal for aversive learning. Mol Psychiatry 27, 3479–3492 (2022).

52. Lan JY, et al. Protein kinase C modulates NMDA receptor trafficking and gating. Nat Neurosci 4, 382–390 (2001).

53. Maekawa M, et al. Signaling from Rho to the actin cytoskeleton through protein kinases ROCK and LIM-kinase. Science 285, 895–898 (1999).

54. Nakahata Y, Yasuda R. Plasticity of Spine Structure: Local Signaling, Translation and Cytoskeletal Reorganization. Front Synaptic Neurosci 10, 29 (2018).

55. Rasmussen AH, Rasmussen HB, Silahtaroglu A. The DLGAP family: neuronal expression, function and role in brain disorders. Mol Brain 10, 43 (2017).

56. Kawano Y, et al. Phosphorylation of myosin-binding subunit (MBS) of myosin phosphatase by Rho-kinase in vivo. J Cell Biol 147, 1023–1038 (1999).

57. Takano T, et al. Discovery of long-range inhibitory signaling to ensure single axon formation. Nat Commun 8, 33 (2017).

58. Uchino S, et al. Direct interaction of post-synaptic density-95/Dlg/ZO-1 domain-containing synaptic molecule Shank3 with GluR1 alpha-amino-3-hydroxy-5-methyl-4-isoxazole propionic acid receptor. J Neurochem 97, 1203–1214 (2006).

59. Amano M, Chihara K, Nakamura N, Kaneko T, Matsuura Y, Kaibuchi K. The COOH terminus of Rho-kinase negatively regulates rho-kinase activity. J Biol Chem 274, 32418–32424 (1999).

60. Hayashi Y, et al. Regulation of neuronal nitric-oxide synthase by calmodulin kinases. J Biol Chem 274, 20597–20602 (1999).

61. Komeima K, Hayashi Y, Naito Y, Watanabe Y. Inhibition of neuronal nitric-oxide synthase by calcium/ calmodulin-dependent protein kinase IIalpha through Ser847 phosphorylation in NG108-15 neuronal cells. J Biol Chem 275, 28139–28143 (2000).

62. Zhang X, et al. Balance between dopamine and adenosine signals regulates the PKA/Rap1 pathway in striatal medium spiny neurons. Neurochem Int 122, 8–18 (2019).

63. Lin YH, et al. Accumbal D2R-medium spiny neurons regulate aversive behaviors through PKA-Rap1 pathway. Neurochem Int 143, 104935 (2021).

64. Amano M, et al. Kinase-interacting substrate screening is a novel method to identify kinase substrates. J Cell Biol 209, 895–912 (2015).

65. Mori K, et al. Rho-kinase contributes to sustained RhoA activation through phosphorylation of p190A RhoGAP. J Biol Chem 284, 5067–5076 (2009).

66. Sooksawate T, et al. Viral vector-mediated selective and reversible blockade of the pathway for visual orienting in mice. Front Neural Circuits 7, 162 (2013).

67. Dickstein DL, et al. Automatic Dendritic Spine Quantification from Confocal Data with Neurolucida 360. Curr Protoc Neurosci 77, 1 27 21–21 27 21 (2016).

68. Surmeier DJ, Ding J, Day M, Wang Z, Shen W. D1 and D2 dopamine-receptor modulation of striatal glutamatergic signaling in striatal medium spiny neurons. Trends Neurosci 30, 228–235 (2007).

